# Mismatch Negativity and General Cognitive Ability - A Meta-Analysis

**DOI:** 10.64898/2026.08.06.743003

**Authors:** Tobias Nöth, Matthew J. Euler, Kirsten Hilger

## Abstract

Intelligence or general cognitive ability (GCA) is fundamental to human behavior and cognition. It impacts important life outcomes like educational success and even health and longevity have been related to differences in GCA. Understanding the biological basis of such individual variations presents a crucial goal of neuroscience. The mismatch negativity (MMN) is an event-related brain potential that can be measured when—within a series of frequent standard stimuli, rare deviants are presented—and is suggested to reflect conscious (active) or unconscious (passive) detection processes of the brain. Importantly, variations in MMN amplitude and latency have been linked to differences in GCA. Yet findings vary considerably. This preregistered meta-analysis provides a comprehensive and structured overview of the current state of research. Following the study selection process in accordance with the PRISMA guidelines, and the rating of study design quality with the Study Design and Implementation Assessment Device for Individual Difference Research (DIAD-ID), the association between GCA and MMN amplitude and latency was examined in 695 healthy adults across 13 included studies. The estimated across-sample associations between MMN and GCA were small, but significant (MMN amplitude–GCA: *r* = −0.08, MMN latency–GCA: *r* = −0.13; *p* < 0.05) and moderators were identified. Between-sample heterogeneity was moderate, with no evidence of publication bias. Our findings suggest that higher cognitive ability is associated with slightly stronger and faster MMN responses. However, the low estimated across-sample effect sizes and the small number of included studies also highlight the need for more research.

## 1. Introduction

Intelligence is a multidimensional construct of general cognitive ability (GCA) that includes the understanding of elaborate concepts, adaptation to environmental changes, problem-solving, and abstract reasoning (Neisser et al., 1996). To quantify individual variations in this complex construct, researchers have developed various intelligence tests with proven reliabilities like the Wechsler Abbreviated Scale of Intelligence (WASI; Wechsler, 1999), the Multidimensional Aptitude Battery (MAB; Jackson, 1984), or the Raven’s Advanced Progressive Matrices (RAPM; Raven, 1965). These measures provide a standardized estimate of individual cognitive capacities, typically expressed as the intelligence quotient (IQ), and are employed in multiple areas, such as in diagnostics (e.g., for learning disabilities or developmental disorders) or educational placement (deLeyer-Tiarks et al., 2024). Beyond these practical applications, GCA has also been linked to long-term health benefits. For instance, higher psychometric childhood intelligence scores have been related to prolonged functional independence and a lower risk of premature mortality (Deary et al., 2004), while cognitive reserve-stimulating behavior has been associated with a reduced vulnerability to the occurrence of dementia (Wang et al., 2017). However, intelligence tests offer limited insight into the underlying mechanisms and factors shaping intelligence across life (Kovacs & Conway, 2019). Investigating the neurobiological foundation of general cognitive ability presents a crucial opportunity to close this gap.

For example, event-related potential (ERP) has been explored as a promising neurobiological indicator of cognitive abilities (Jausovec & Jausovec, 2000): ERPs are small electric currents that are recorded on the scalp and reflect neurological brain processes evoked by (sensory) input (Blackwood & Muir, 1990) or other experimental events. These electric signals can be measured by using the non-invasive technique of electroencephalography (EEG), which offers higher temporal resolution compared to other neuroscientific technologies like functional magnetic resonance imaging (fMRI) or positron emission tomography (PET) (Cao et al., 2021; Yen et al., 2023). This fine temporal sensitivity allows dividing ERPs into smaller components that differ on a millisecond (ms) timescale (Luck, 2014), while reflecting different neuronal processes. ERPs are characterized by a positive peak (e.g., the P3) or a negative peak (e.g., the N100) following stimulus onset, as well as by their waveform (Sur & Sinha, 2009). Some specific components are calculated by subtracting the ERP of one experimental condition from the ERP of another condition (so-called difference waves, e.g., the Mismatch Negativity, MMN). Crucially, individual variations in ERP components have been linked to differences in GCA (Hilger et al., 2022).

While the association between GCA and the P3 (also referred to as P300) component has been intensively researched throughout the last decades (for review see Hilger et al., 2022), findings have been highly heterogeneous and often even contradicting each other (Li et al., 2021; Russo et al., 2008; Walhovd et al., 2005; Walhovd & Fjell, 2003). A recent meta-analysis addresses this heterogeneity and reports a significant but small positive across-study association between GCA and P3 amplitude of *r* = 0.13 as well as a significant negative across-study association of *r* = −0.18 between GCA and P3 latency (Euler & Hilger, 2026). Another ERP component that has been linked to the biological foundation of GCA is the MMN. The MMN can be obtained in response to an infrequent and unexpected deviant stimulus (Näätänen et al., 1978) and has been classified as part of the N2 component, i.e., the second large negative deflection in the ERP waveform that peaks around 200ms after stimulus onset (Näätänen & Gaillard, 1983). In previous research, the N2 component has been associated with cognitive control and its mechanisms (Folstein & van Petten, 2008), posing the question of which role MMN plays in regulatory executive mechanisms relevant to GCA.

Typically, the MMN is assessed using so-called passive auditory oddball paradigms (O’Reilly & O’Reilly, 2021; Czigler & Kojouharova, 2022). These tasks are usually composed of an active component (e.g., reading a book, a simple change detection task) occupying the participant’s attention and a passive oddball component, involving the presentation of a frequent, recurring standard stimulus and occasionally rare deviant stimuli (the oddballs). Both stimuli are passively perceived and processed by the subject, with the oddball stimulus evoking the MMN. Once the MMN is elicited, two primary measures are typically assessed in the MMN time window (100-250ms after stimulus onset): MMN peak amplitude, reflecting the highest peak of the difference wave in that time frame (calculated as the ERP elicited by deviant stimuli minus the ERP elicited by standard stimuli; Fitzgerald & Todd, 2020), and MMN peak latency, referring to the time between stimulus onset and the highest peak of that difference wave (Duncan et al., 2009). A well-established theory, in line with basic theories concerning the generation of MMN, links MMN to cognitive ability (Näätänen et al., 2007): The authors argue that the MMN reflects underlying mental processes in the auditory cortex, potentially indicative of some form of “primitive sensory intelligence”. However, the present state of evidence in this field is heterogeneous. While many studies report associations between increased MMN amplitudes and psychometric intelligence, the exact sizes of these correlations vary considerably (ranging from −0.31 to 0.41, e.g., Beauchamp & Stelmack, 2006; De Pascalis & Varriale, 2012; De Pascalis et al., 2014), and a recent study employing a visual MMN paradigm found no evidence of an association between MMN amplitude and cognitive ability at all (Hilger & Euler, 2023). Concerning the relation between GCA and MMN latencies, the picture is similar. Some studies report strong negative associations up to *r* = −0.61 (Beauchamp & Stelmack, 2006), while others found mixed results (Hilger & Euler, 2023) or even weak positive associations (*r* = 0.11, Getzmann et al., 2013). This inter-study inconsistency highlights the need for a structured, comprehensive overview to ultimately achieve a deeper understanding of the neurobiological mechanisms of general cognitive ability.

The main goal of this meta-analysis is to address this gap by providing a systematic comparative review of the relation between the mismatch negativity and general cognitive ability. Specifically, this study compares the association between MMN latency and amplitude and GCA over 13 scientific articles. These papers have been selected after a comprehensive and systematic search and screening process in accordance with the PRISMA guidelines. Methodological quality and risk of bias in the included studies were evaluated with the adapted version of the Study Design and Implementation Device (DIAD, Valentine & Cooper, 2008; DIAD-ID, Euler & Hilger, 2026), and various moderators were explored. Finally, potential reasons for the observed heterogeneity, critical limitations, publication bias, and future perspectives in this field of research were discussed.

## 2. Hypotheses

This meta-analysis assesses the potential relationship between the amplitude of MMN and GCA, as well as between the latency of MMN and GCA, while also exploring potential moderators of these associations. To this aim, this study will focus on two specific hypotheses that were derived from the available empirical evidence and have been preregistered on the Open Science Framework (OSF): https://osf.io/59n6u.

**H1.** There exists a significant negative association between the amplitude of the mismatch negativity and general cognitive ability across studies (e.g., Houlihan & Stelmack, 2012).

**H2.** There exists a significant negative association between the latency of the mismatch negativity and general cognitive ability across studies (e.g., De Pascalis et al., 2014).

Furthermore, the influence of the following potential moderators on the association between MMN amplitude and latency and GCA will be explored: task difficulty, mean age, gender, and EEG recording window length.

## 3. Method

### 3.1 Preregistration

The hypotheses and all methodological details of this study were formally preregistered in the Open Science Framework: https://osf.io/59n6u. Note, however, that during the literature search phase, unanticipated complications led to some modifications to the preregistered search string for specific databases. These were necessary to match the specifications of these databases while still adhering to the preregistered plan as closely as possible. Furthermore, during title and abstract screening and the full-text review stage, the need to specify more specific exclusion criteria, as well as specific criteria for measures of general cognitive ability, arose. These changes were made to ensure comparative validity between the included studies and are described in further detail in the respective sections (3.2, 3.2.1, 3.2.2).

### 3.2 Literature Search and Study Selection

The literature search was carried out between March and April of 2025; only studies published before that time were considered in this meta-analysis. To determine the best-suited search string that would yield the most sufficient results, multiple pilot test searches were conducted. Focusing on non-clinical peer-reviewed articles, the following databases were employed: PubMed, Google Scholar, APA PsycNet, Scopus, and Web of Science. After probing the search query on these databases, the final search string was defined as: (MMN OR Mismatch negativity OR mismatch-negativity) AND (“cognitive ability” OR intel* OR “IQ” OR “general intelligence” or “fluid intelligence”) AND (ERP OR “event-related potential” OR EEG).

However, while trialing the search string, it already became clear that modifications to it would be necessary for some databases due to varying requirements as well as different search mechanisms. The literature search on PubMed, Google Scholar, and APA PsycNet could be carried out without altering the string, yielding 149 results for PubMed, 232 results for Google Scholar, and 223 for APA PsycNet. Due to limitations regarding the length of the search string on Scopus, the search query was adjusted. The following search inquiry allowed a comparable outcome by removing parts of the search string, resulting in a slightly broader scope and thus ensuring that no potentially relevant study would be overlooked: (MMN OR Mismatch negativity OR mismatch-negativity) AND (“cognitive ability” OR intel* OR “IQ” OR “general intelligence” or “fluid intelligence”). This search resulted in 227 studies. Furthermore, during the time of the literature search, the filter tool of Web of Science did not execute the “AND” function as intended, leading to over 160000 results with the original query. Thus, this search procedure was also altered: The search string was split into its three main components, and these were then applied in the following order: (MMN OR Mismatch negativity OR mismatch-negativity), then (“cognitive ability” OR intel* OR “IQ” OR “general intelligence” or “fluid intelligence”) and after that (ERP OR “event-related potential” OR EEG). This filtered the results of each search by the next query component, ultimately yielding 165 results. Finally, one additional paper (Jäncke et al., 2012) was identified through the author contact during the full-text review phase. This study was subsequently included and subjected to all stages of the meta-analysis process.

#### 3.2.1 Title and Abstract Screening

The literature search culminated in a total of 997 articles, which were first processed via the Zotero research assistant tool (https://www.zotero.org) and then uploaded to the Covidence systematic review management software for screening (https://www.covidence.org). This software automatically removes duplicates from the uploaded articles, resulting in 276 redundant entries being removed and leaving a total of 721 studies for the next step, the title and abstract screening (Figure 1). This first screening stage was independently conducted by two reviewers, who rejected any articles that matched any of the following exclusion criteria: use of non-human subjects; focus on children or adolescents (< 18 years); primary focus on neurological, psychiatric, or medical conditions (e.g., autoimmune disorders, endocrine/metabolic conditions, etc.); MMN was not measured from scalp electrodes or measurement approach was too idiosyncratic to allow coding of electrodes, measurement windows, etc.; article not available in English or German. Articles that did not reference individual differences in cognitive processes, tasks, or assessments in the abstract or articles that could not be classified as an empirical study (e.g., meta-analyses, reviews, books or book chapters, commentaries, etc.), were also excluded. If the information necessary to make a conclusive decision about the exclusion of an article could not be derived from the title and abstract, the study was moved to the full-text review stage and assessed again. Articles focused on interventions (e.g., training programs aiming to enhance cognitive abilities) were advanced to the next stage. Conflicts between the ratings of both reviewers were discussed and resolved, leaving 272 studies for the full-text review.

**Figure 1.**
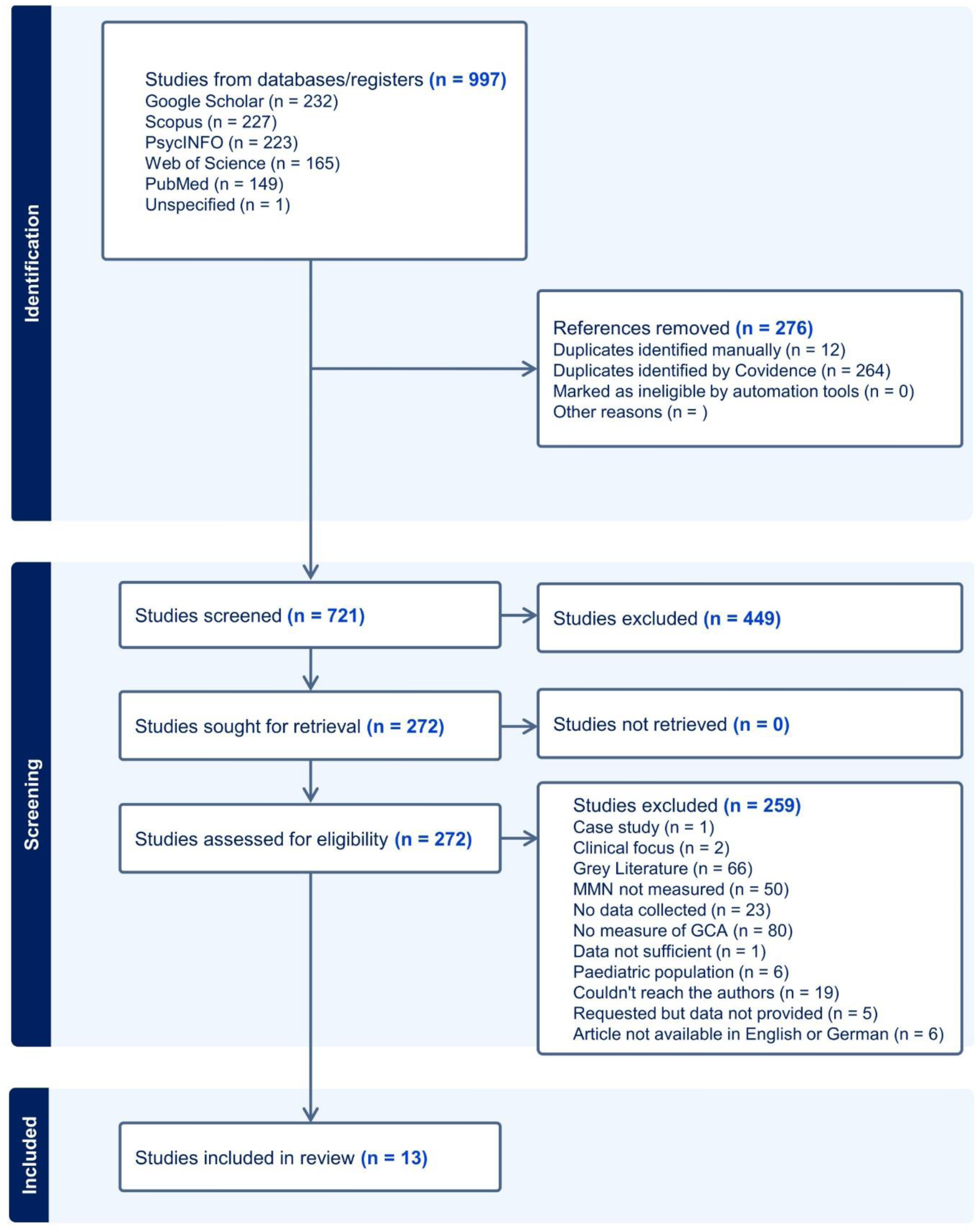
Flowchart showing the study selection process based on the PRISMA guidelines. Of 997 identified articles, 13 qualified for inclusion in the qualitative review and quantitative meta-analysis.

#### 3.2.2 Full-Text Review

During the full-text review stage, two independent reviewers evaluated the remaining articles, while considering two additional exclusion criteria at this phase of the meta-analysis: a) the study did not report any correlation value for the association between MMN and GCA, nor the average MMN amplitude/latency values in studies comparing higher vs. lower ability subjects and, b) non-peer-reviewed gray literature (e.g., posters, theses, preprints, books and book chapters, etc.). Furthermore, in order to define GCA more precisely, another un-preregistered exclusion criterion was added by specifying the measure of GCA as the following: c) GCA had to be measured via a standardized, published and established test of GCA (e.g., the Multidimensional Aptitude Battery, Jackson, 1984; the Raven’s Advanced Progressive Matrices, Raven, 1965; and the Wechsler Adult Intelligence Scales and subtests, Wechsler, 1955) or via a subtest of these measures. Additionally, studies that used tests that are mainly focused on assessment of cognitive and neuropsychological functioning as well as on clinical diagnoses (e.g., word-list learning tasks, the Wisconsin Card Sorting Task; Sherman et al., 2023) were excluded. However, owing to their strong correlation with IQ and to the established use as substitute indices of GCA, validated and standardized tests of irregular word reading like the National Adult Reading Test (NART; Nelson, 1982) were also accepted as measures of GCA (Blair & Spreen, 1989).

During this stage, the previous exclusion criteria were assessed again, and studies that did not match any exclusion criteria and reported a correlational analysis between at least one measure of MMN and an adequate measure of GCA were moved on to the data extraction phase. In cases where a study did not meet any of the exclusion criteria except for not reporting a correlation between MMN and GCA, the authors of the article were contacted via email and asked whether they would be able to provide a correlation or sufficient data to calculate the correlations (amplitude, latency, GCA measures), while outlining a clear response deadline. As before, all contrasting evaluations between the two reviewers were addressed and resolved. Finally, a total of 13 studies qualified for the extraction phase and were thus included in this systematic meta-analysis.

### 3.3 Data Extraction and Variables of Interest

The following information was extracted from each study:

- <u>MMN-GCA Correlation:</u> Primary variable of interest: any association between MMN amplitude or MMN latency and a measure of GCA, as reported in the original study text, directly supplied by the study authors, or computed out of the data provided by the authors.
- <u>General Study Information:</u> DOI, names of the authors, publication year, and whether the data were reported in the original study text or provided by the authors.
- <u>Sample Information:</u> Original description of the study sample (e.g., “English-speaking women”, “female psychology students”, etc.), final sample sizes (after any removals, for more details, see “*Procedure for provided data”*) were documented for each MMN-GCA correlation separately, number of subjects below 18 years old, descriptive statistics regarding participants age (minimum (“youngest”), maximum (“oldest”), median, mean and standard deviation), percentage of female, male and non-binary participants, participants’ handedness, study location, educational and occupational demographics.
- <u>GCA:</u> Exact name of the intelligence measure, specific domain supposed to be measured by the test (e.g., MAB: general intelligence; Ravens matrices: fluid intelligence, etc.), amount of (sub)tests applied, testing time, descriptive statistics for intelligence scores (minimum, maximum, mean, standard deviation) and whether GCA scores were reported in the text or provided by the authors.
- <u>Task Characteristics:</u> Name of the task chosen by the authors of the original study, classification of task type by the authors of this meta-analysis (Oddball, Attention or Cognitive Control, Working Memory, and Reaction Time/Chronometric). If no clear classification was possible, the tasks were consolidated into a single category “Other”. Oddball tasks were categorized as either active or passive, as well as by stimulus type (Standard, Target, Novelty, and Other). Further, it was extracted whether the task inter-stimulus interval was fixed or not.
- <u>Task Difficulty:</u> Some studies reported multiple MMN-GCA correlations originating from different conditions within a task, such as changes in deviant frequency (e.g., Bazana & Stelmack, 2002), pitch (Jäncke et al., 2012), or changes to the temporal characteristics of the stimulus (Troche et al., 2010). For these studies, difficulty was estimated based on the stimulus differences (smaller differences between very similar stimuli were coded as harder to detect and thus as more difficult) or on the difficulty degrees implied in respective text passages from the original article. In case this information was inconclusive, difficulty was not coded.
- <u>EEG Recording:</u> Final sampling rate, online and offline reference, online and offline high-pass, low-pass, and notch filters.
- <u>MMN:</u> specific type of MMN latency and amplitude measure used (e.g., peak, signed area, etc.), amplitude and latency time windows (start and end time in milliseconds), descriptive statistics (mean, standard deviation, minimum, and maximum) for MMN amplitude and latencies at the electrodes Fz, FCz, Cz, Pz, POz, Oz, and “Other”.

Note that since some studies consisted of several subsamples (due to sub-studies and sub-groups or multiple tasks within one study), MMN-GCA correlations were coded per sample whenever possible, and information for each sample was extracted separately. All following analyses were thus conducted at the sample level rather than the study level.

Exceptional Cases and Caveats: Due to some studies only reporting demographic characteristics for the full sample (without providing detailed information for subsamples or for the finally employed sample, e.g., after outlier exclusion took place), only demographic data that could be clearly attributed to the relevant subsamples were included in the meta-analysis. Furthermore, for studies that reported mean IQ values across a low- and a high-ability group of the same size, the average IQ score for the whole sample was calculated.

*Procedure for Provided Data:* For studies that did not directly report correlations but provided the necessary data, correlations were calculated via Spearman’s *rho*. This was decided after assessments of the data regarding linearity, normality, homoscedasticity, and absence of outliers showed that the data did not fulfill the assumptions required to calculate Pearson’s *r*. Within the provided datasets, any individual values straying more than three standard deviations from the mean were removed. Furthermore, given that the MMN is a negative component, derived by removing the elicited activity of the standard stimulus from the activity elicited by the deviant (Fitzgerald & Todd, 2020), participants exhibiting positive MMN amplitudes were excluded from the analysis examining the association between MMN amplitude and general cognitive ability. More details about the included samples are provided in the online Supplementary Materials. After applying these steps, the correlations between MMN amplitude/latency and the respective GCA measures were calculated and entered into the extraction sheet along with the other relevant information mentioned above.

### 3.4 Study Quality Design Assessment

To provide a systematic and objective estimate of study quality and to evaluate potential bias in study design, all included studies were assessed with the Study Design and Implementation Assessment Device for Individual Difference Research (DIAD-ID; Euler & Hilger, 2026), which was adjusted to the specific parameters and characteristics of the MMN for the present meta-analysis. The DIAD-ID is constructed as a system of 11 overarching contextual questions covering construct, external, and statistical validity (for more details, see Euler & Hilger, 2026). It presents a further development of the Study DIAD by Valentine & Cooper (2008) that primarily focused on intervention studies and has been used extensively (Bernard et al., 2014; Linhardt et al., 2022; Pfeiffer et al., 2024). To ensure comparability and transparency, the adapted DIAD-ID, including adjusted rating categories and study ratings, has been made available online: https://osf.io/59n6u

### 3.5 Examination of Outliers

The outlier analysis at the single study level consisted of the combination of a Cook’s distance value larger than the median plus six times the interquartile range and of studentized residuals ± 1.96 (Viechtbauer & Cheung, 2010). All associations between GCA and MMN amplitude, as well as between GCA and MMN latency, were assessed using these criteria (Pfeiffer et al., 2024). This identified two samples, of which neither met the joint criterion. Nonetheless, to estimate their general impact on meta-analytical across-sample associations, sensitivity analyses excluding these two studies were carried out.

### 3.6 Meta-Analytic Synthesis of Results

All analyses were carried out in R (version 4.5.2; R Core Team, 2025) using the following packages: tidyverse (Wickham et al., 2019), RColorBrewer (Neuwirth, 2022), moments (Komsta & Novomestky, 2022), pander (Daroczi & Tsegelskyi, 2025), psych (Revelle, 2025), stringr (Wickham, 2025), forestplot (Gordon & Lumley, 2025), metafor (Viechtbauer, 2010), purr (Wickham & Henry, 2025) and readxl (Wickham & Bryan, 2025). Three additional packages were used for testing the statistical assumptions of the provided data: olsrr (Hebbali, 2024), lmtest (Zeileis & Hothorn, 2002), and patchwork (Pedersen, 2025). The R-code was based on the code of Euler and Hilger (2026) and adapted to fit the requirements of the MMN. First, the correlations between MMN amplitude and latency (respectively) and GCA were Fisher’s *z*-transformed and subsequently weighted by sample size. To evaluate the two main hypotheses regarding the associations between GCA and MMN amplitudes (H1) and latencies (H2), intercept-only meta-analytic models were applied, with sample specified as a random effect. This method provided the opportunity to compute the mean effect size for amplitude and latency associations across samples with between-sample heterogeneity explicitly considered. The potential moderating role of task difficulty on the association between GCA and MMN amplitude and latency was tested by setting difficulty as a categorical fixed effect with three levels (“Low”, “Medium”, “High”), with Low as the reference category. Sample was again incorporated as a random effect. The influences of the continuous moderators, such as mean age, gender (quantified by the percentage of females), and window length, on the associations between GCA and MMN measures were examined using separate multilevel meta-regression analyses for each moderator with random intercepts for study, subsample, and effect. All moderators were mean-centered prior to analysis by subtracting their study-level mean. Once statistical assumptions were verified, model fitting was performed using the Restricted Maximum Likelihood estimator (default in metafor). To control for Type I errors in moderation analyses, *p*-values were Bonferroni corrected (Armstrong, 2014).

### 3.7 Publication Bias

Publication bias was assessed across samples. First, Fisher’s *z* transformation and fixed-effect inverse-variance weighting were performed, and samples with more than one correlation were aggregated. Additional steps included visual inspection of graphic funnel plots and analyses for statistical asymmetry. The latter was performed by applying Egger’s regression (Egger et al., 1997) as well as the rank correlation test (Begg and Mazumdar, 1994) to a univariate random-effects model. When these analyses indicated asymmetry, trim-and-fill imputation was employed to estimate adjusted correlations. To gauge the potential influence of the file-drawer problem, the generalized fail-safe *N* (Rosenthal, 1979) was computed, providing an approximation of the number of studies necessary to change the meta-analytic overall effect to non-significance with alpha = .05.

### 3.8 Data and Code Availability Statement

This meta-analysis relied on open-source software and packages. All code used for calculations and to generate the figures in this paper, the adapted and blank DIAD-ID template, the completed DIAD-ID, as well as the full extraction database, including the final extraction sheet, are accessible online via OSF: https://osf.io/59n6u.

## 4. Results

### 4.1 Systematic Review

*4.1.1 Study Sample Characteristics*

After exclusion, 13 studies were eligible for quantitative meta-analyses, amounting to 14 unique samples, *N* = 695 participants, and 158 total MMN-GCA correlations (including all associations for both amplitudes and latencies across samples and subsamples, varying experimental conditions and electrodes). The number of correlations reported per study varied, ranging from 2 to 36 (reported data: 4-36; provided data: 2-16; Table 2) and sample sizes also differed widely (Mean (M) = 49.26; Standard Deviation (SD) = 36.12; range: 21-160) revealing that most studies did not meet recommended power levels for individual difference research (Brysbaert, 2019; DeYoung et al., 2025; Gignac & Szodorai, 2016). The typical study sample predominantly contained right-handed women in their early to mid-twenties. However, five out of the seven studies reporting minimum and maximum ages for the final sample included at least one subject over 40, with one study covering a range from 18 to 65+ (Berti et al., 2012). In one of the studies included in this meta-analysis, only participants between 34 and 56 were examined (Bishop et al., 2011), presenting an exception to the typically student-heavy research populations. While pediatric samples were excluded and underage participants were removed from the provided data, one study included at least one participant under 18 years of age (Troche et al, 2010). The professional background of the participants was reported in eight studies, with six studies including student-only samples. The other two studies focused (partially) on professionals in their respective fields (Jäncke et al., 2012; Rogenmoser et al., 2015). Educational level was not reported in any of the studies included.

**Table 1.**
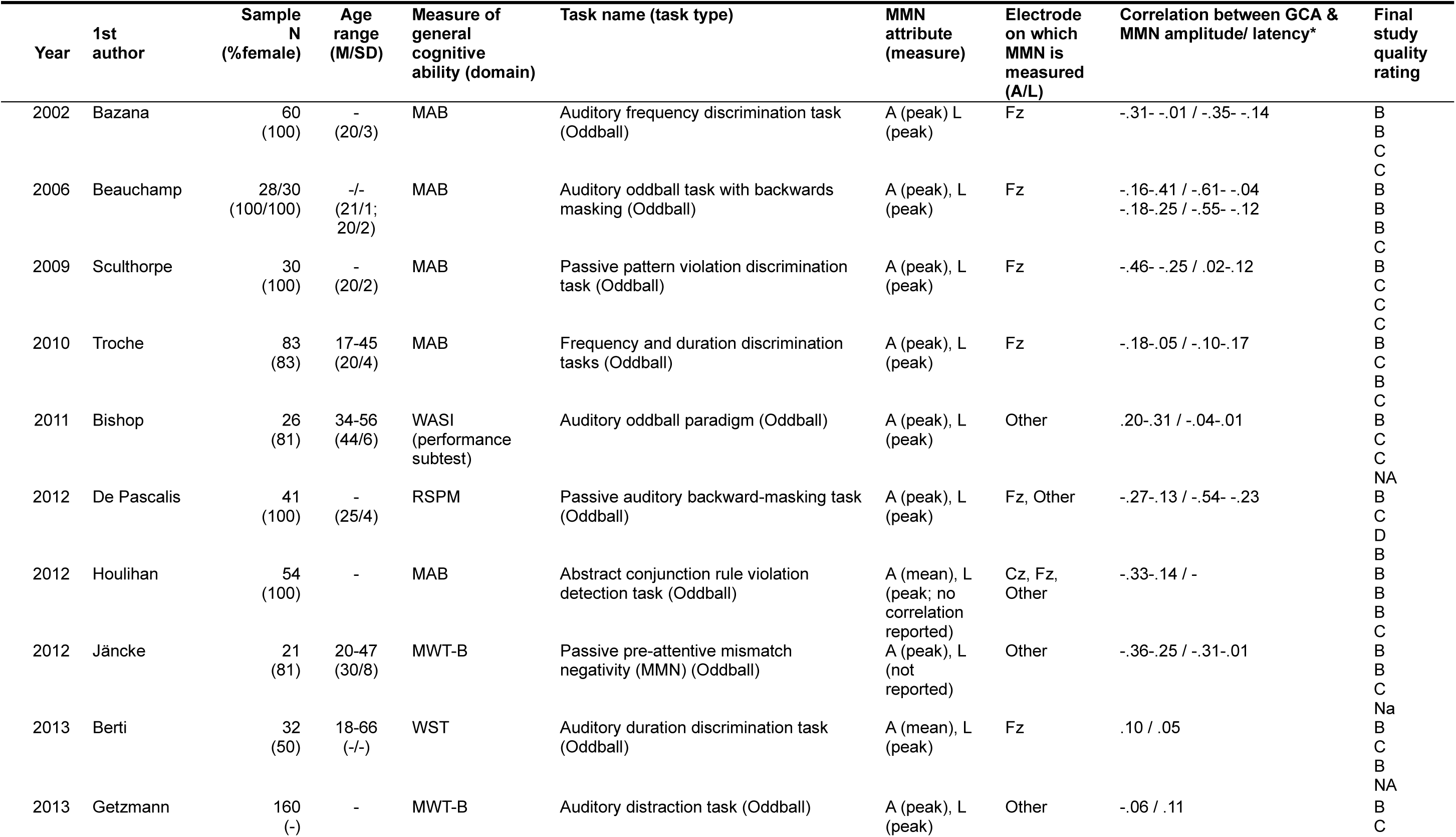
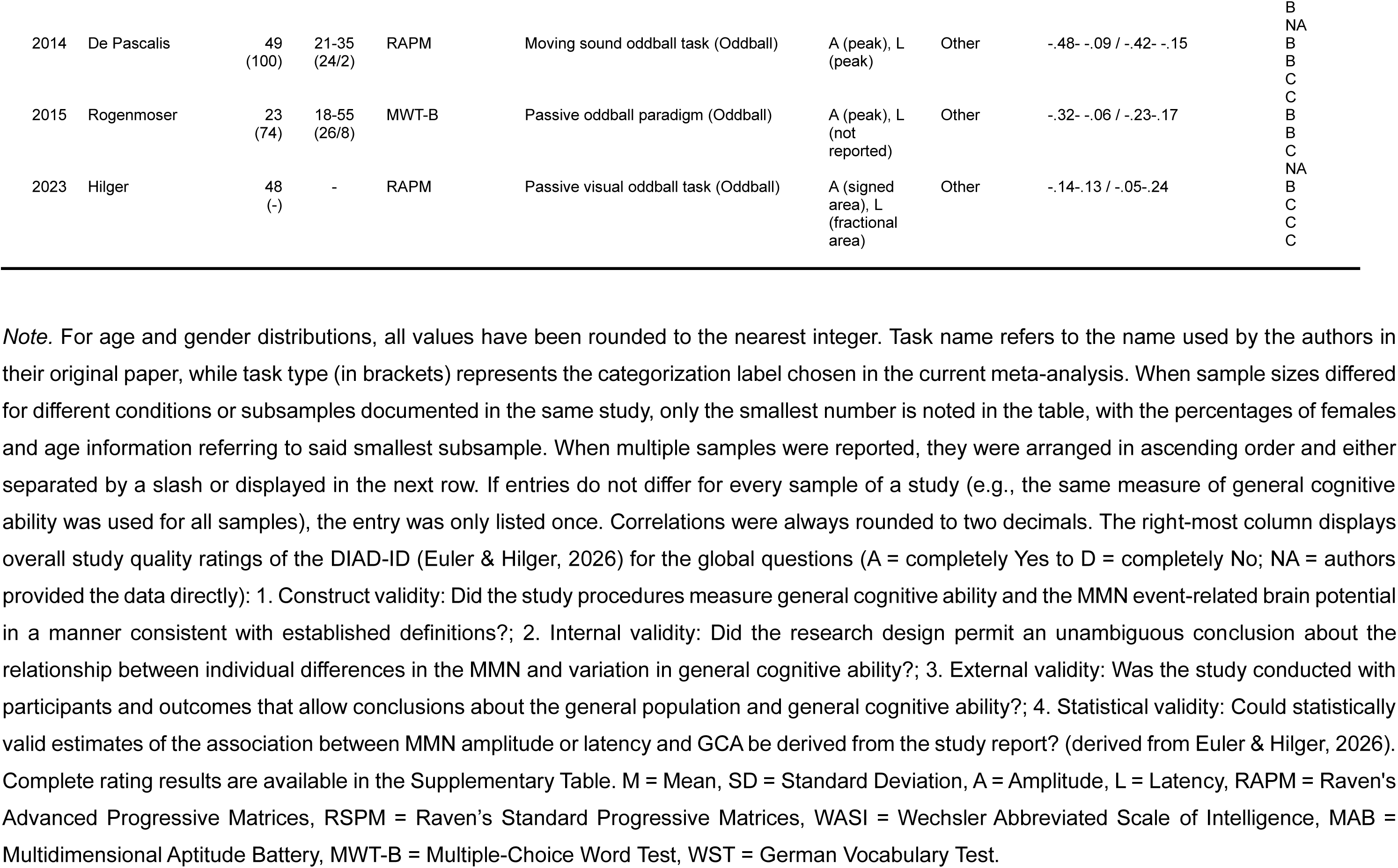
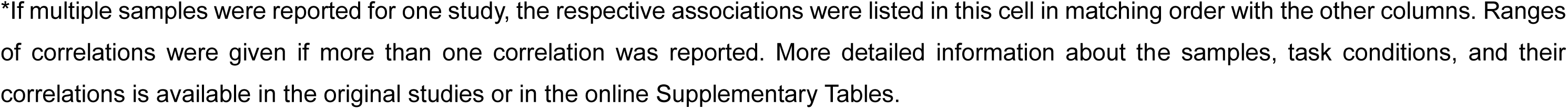
Studies included in the meta-analyses in chronological and alphabetical order.

**Table 2.** Demographic characteristics of the meta-analytic sample and reporting frequencies.

| *Variable* | *Mean* | *Median* | *SD* | *Min* | *Max* | *Skew* | *Reporting Frequencies* |  |
| --- | --- | --- | --- | --- | --- | --- | --- | --- |
|  |  |  |  |  |  |  | *Studies* | *Samples* |
| Sample Size | 49.26 | 36.5 | 36.12 | 21.5 | 160 | 2.24 | 13 | 14 |
| Mean Age | 24.79 | 23.64 | 7.07 | 19.5 | 43.67 | 1.89 | 10 | 11 |
| SD Age | 3.81 | 3.62 | 2.09 | 1.2 | 7.69 | 0.64 | 10 | 11 |
| Age Youngest | 21 | 18 | 5.94 | 17 | 34 | 1.76 | 7 | 7 |
| Age Oldest | 46 | 45 | 11.54 | 33 | 66 | 0.52 | 7 | 7 |
| % Female | 88.03 | 100 | 15.62 | 50 | 100 | -1.11 | 12 | 13 |
| % Right-Handed | 99.36 | 100 | 1.70 | 95.5 | 100 | -2.04 | 6 | 7 |
| Sample IQ Mean | 111.7 | 114.1 | 7.19 | 92.32 | 117.4 | -2.20 | 9 | 10 |
| Sample IQ SD | 13.97 | 12.11 | 4.86 | 10.3 | 23.3 | 1.38 | 6 | 6 |
| Sample Minimum IQ | 81.4 | 87 | 17.47 | 61 | 100 | -0.22 | 5 | 5 |
| Sample Maximum IQ | 137 | 141 | 8.12 | 123 | 143 | -1.25 | 5 | 5 |

*Note.* All values have been rounded to the nearest decimal. Studies = number of studies reporting the variable of interest. Samples = number of samples for which that variable is reported. The minimum sample size value is the average of the subsamples within one study (Jäncke et al., 2012), since two participants had to be removed for some parts of the analysis. % = percentage. IQ = intelligence quotient.

#### 4.1.2 Cognitive Ability Measures

General intelligence (42.9%), fluid intelligence (28.6%), and crystallized intelligence (28.6%; percentages refer to samples of included studies, Figure 2) were all investigated for their association with MMN amplitudes/latencies. The mean number of tests administered to measure cognitive ability for each sample was 4.92, with a median (MD) of 1.5 (SD = 4.57; range 1-10). These tests covered on average one dimension of the broader GCA construct (e.g., fluid intelligence or crystallized intelligence; M = 1.43; SD = 0.51; MD = 1; range: 1-2).

**Figure 2.**
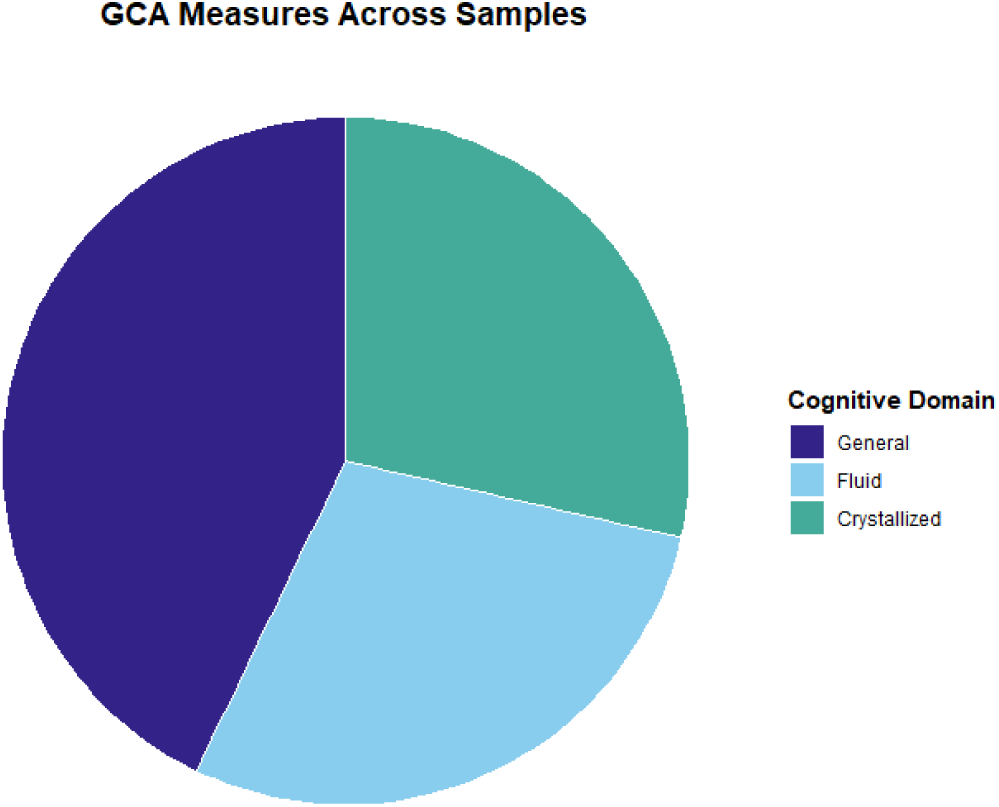
The assessed general cognitive ability domains were balanced between general, fluid, and crystallized intelligence. Proportions of reported domains refer to studies included in the meta-analysis.

For ten samples, mean standardized IQ values were reported, resulting in an average mean IQ of 111.7 (SD = 7.19; range: 92.32-117.4; skew = −2.20). Additionally, SDs for the mean standardized IQ values were documented for six of these samples, averaging to an across-sample IQ SD of 13.97 (M; range of SDs: 10.3-23.3). Within the five studies reporting IQ ranges, the lowest and the highest IQ scores per sample varied broadly, ranging from 61 to 100 (Minimum IQ; M = 81.4) and from 123 to 143 (Maximum IQ; M = 137). To summarize, the study samples included in this meta-analysis are characterized by above average IQ values, which is also reflected in the negative skew of the IQ score distribution, and a nearly similar, but slightly smaller variation around the mean than the general population.

#### 4.1.3 Study Task Characteristics

The MMN was always elicited by variations of the passive oddball paradigm (100%) with twelve studies employing an auditory task (92.3%) and one study implementing a visual task (7.7%; Hilger & Euler, 2023).

#### 4.1.4 EEG Methodology and Signal Processing

Among the reported recording parameters and preprocessing steps, digital sampling rates were documented in 12 of the 13 included studies (92.3%), with the most common rate being 1000 Hertz (Hz; Mode; Range: 250-1000). Online high-pass and low-pass filters were reported six times (46.2%) and 11 times (84.6%), respectively, with modal values of 0.1 Hz (high-pass filter; Range: 0-0.1 Hz) and 100 Hz (low-pass filter; Range: 30-280 Hz). For offline high- and low-pass filters, the information was reported nine times (high-pass filter; 69.2%) and 12 times (low-pass filter; 92.3%), with modal values of 1 Hz (high-pass filter; Range: 0.05-3 Hz) and 20 Hz (low-pass filter; Range: 10-30 Hz). Three studies (23.1%) reported applying an online notch filter, which were all set to 50 Hz, while no notch filters were applied offline. The most common location used for online referencing was the nose, which was used eight times (61.5%). Other choices included the linked ears (n = 2; 15.4%), FCz (n =1; 7.7%) and one single mastoid (M1/M2; n = 1; 7.7%). One study did not report online references (7.7%). Offline references were only reported sparsely, with 11 studies (84.6%) not reporting an offline reference. Of the two studies (14.4%) reporting offline references, one used linked Mastoids (7.7%) and the other one used the average of all scalp electrodes (7.7%) as references.

#### 4.1.5 ERP-Reporting

Associations for MMN amplitudes were provided in all 13 studies (100%), while 12 (92.3%) additionally reported associations for latencies. Thus, one study reported associations only for MMN amplitudes (Houlihan & Stelmack, 2012). Critically, only five studies (41.7%) reported more detailed descriptives for MMN amplitudes and latencies. Those provided at least amplitude and latency mean scores (one of these studies did not provide the scores for all correlations), with four studies also providing SDs and three also providing minimum and maximum scores. Regardless of the task type deployed during EEG recording, in these five studies, amplitude values recorded at the Fz electrode were descriptively higher than on electrodes labelled as “Other”. The earliest MMN latency peak occurred at the electrodes labelled as “Other”, followed by Fz. However, of the five studies reporting these descriptive data, only two provided values at Fz, with one of them only providing a fraction of the data. Moreover, no data were available for the Pz and Cz electrodes. Thus, only for 19 of the 87 MMN amplitude associations and only for 18 of the 71 MMN latency associations, descriptive data were reported (Table 3), and these descriptive results should be interpreted with caution.

**Table 3.** Descriptive statistics for MMN amplitudes and latencies by electrode.

| *Variable* | *Electrode* | *Mean* | *SD* | *Min* | *Max* | *Samples* | *Effects* | *Total Effects* |
| --- | --- | --- | --- | --- | --- | --- | --- | --- |
| Amplitudes | Fz | -1.28 | 0.54 | -0.9 | -1.66 | 2 | 5 | 37 |
|  | Cz | --- | --- | --- | --- | 0 | 0 | 4 |
|  | Other | -1.12 | 0.74 | -0.31 | -1.76 | 3 | 14 | 46 |
| Latencies | Fz | 184.3 | 12.7 | 175.3 | 193.4 | 2 | 4 | 33 |
|  | Other | 148.4 | 51.1 | 138.9 | 161.0 | 3 | 14 | 38 |

*Note.* Mean, SD, Min, and Max values represent the average values for each statistic across all available samples (e.g., Min = average of the lowest reported amplitude across samples). Samples represents the number of subsamples reporting mean MMN scores, since these were documented most frequently. Effects refers to the number of mean values documented across all samples, task conditions, and electrodes included in this meta-analysis. Total Effects describes the number of associations reported for values from each electrode across all samples. Other refers to any electrode(s) or electrode combination that does not include Fz, Cz, and Pz. It should be noted that Min and Max scores for MMN Amplitudes were sorted by absolute values since larger negative numbers represent larger deflections.

All included studies listed the specific amplitude measure, with 10 studies using peak amplitude (76.9%), two studies using mean amplitude (15.4%; Berti et al., 2013; Houlihan & Stelmack, 2012), and one other study using scalp-averaged signed area amplitudes (7.7%; Hilger & Euler, 2023). Ten of the 12 studies reporting correlations for latency clearly described which latency measurements they deployed. Nine of these studies used peak latency (75%), and one study used fractional area latency (8.3%; Hilger & Euler, 2023). Two studies did not provide clear information about the type of latency measurement they applied (16.7%; Jäncke et al., 2012; Rogenmoser et al., 2015).

The most often implemented time window used to measure MMN was 100-240 milliseconds (ms). This accounted for 21 of the amplitude (24.1%) and for 21 of the latency (29.6%) time windows for MMN and GCA associations. For MMN amplitude, other windows accounting for at least 10% of the associations include 150-150ms (16 correlations; 18.4%), 170-250ms (12 correlations; 13.8%), and 100-250ms (10 correlations; 11.5%). For MMN latency, time frames between 170-250ms (12 correlations; 16.9%) and between 100-250ms (10 correlations; 14.1%) were employed for more than 10% of the assessed correlations. One study did not clearly list the windows used to measure MMN (four correlations for MMN amplitude and MMN latency each; 4.6% and 5.6%, respectively; Jäncke et al., 2012). If the onset and offset times were examined individually, most measurement windows started after 100ms (90 correlations out of 150; 60%) and ended around 240ms (42 correlations; 28%) and 250ms (60 correlations; 40%; for MMN latency and amplitude combined). The earliest onset time was 100ms, and the latest offset time was 400ms post stimulus. For MMN amplitude-GCA associations, the predominant window lengths were 80ms (12 correlations; 14.46%), 100ms (17 correlations; 20.48%), 140ms (21 correlations; 25.30%), and 150ms (14 correlations; 16.87%). Time windows for MMN latency and GCA associations showed a similar trend, with the most common time frames being 80ms (12 correlations; 17.91%), 140ms (21 correlations; 31.34%), and 150ms (14 correlations; 20.90%; Figure 3).

**Figure 3.**
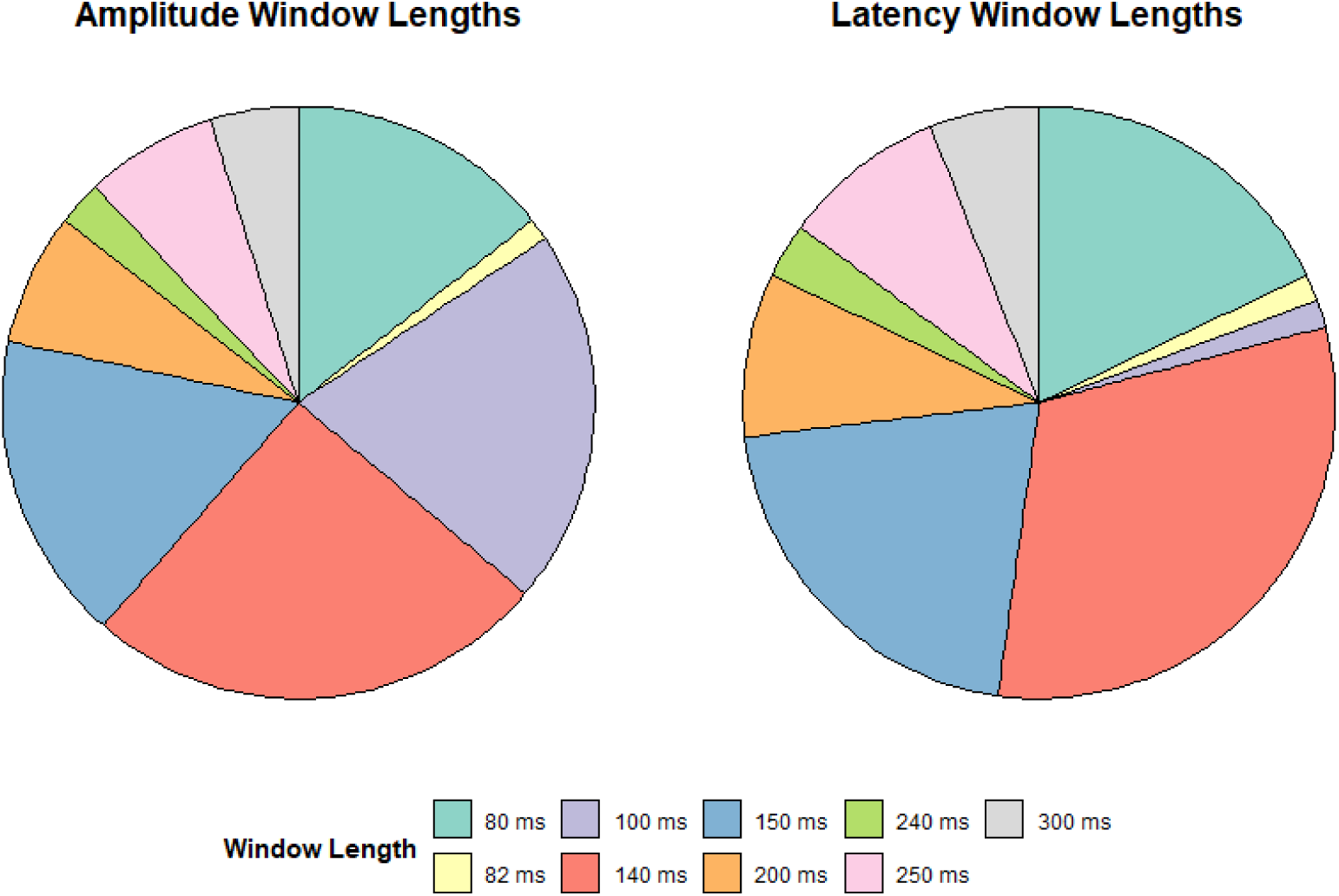
MMN amplitude and latency were measured during time windows of varying size. Amplitude Window Lengths: Size of each segment reflects the relative number of correlations (total: 83 correlations) between MMN amplitude and GCA recorded within the respective time frame. Latency Window Lengths: Size of each segment reflects the relative number of correlations (total: 67 correlations) between MMN Latency and GCA recorded within the respective time frame. Note that one study did not report exact time windows (Jäncke et al., 2012).

### 4.2 Study Quality Assessment (DIAD-ID)

For every study, the respective global DIAD-ID quality score is reported in the rightmost column in Table 1 as well as in Figure 4. An overview of the adapted questions and the rating algorithm, as well as more detailed results regarding design, implementation, and contextual questions, are available in the Supplementary Material on OSF: https://osf.io/59n6u/files/osfstorage.

**Figure 4.**
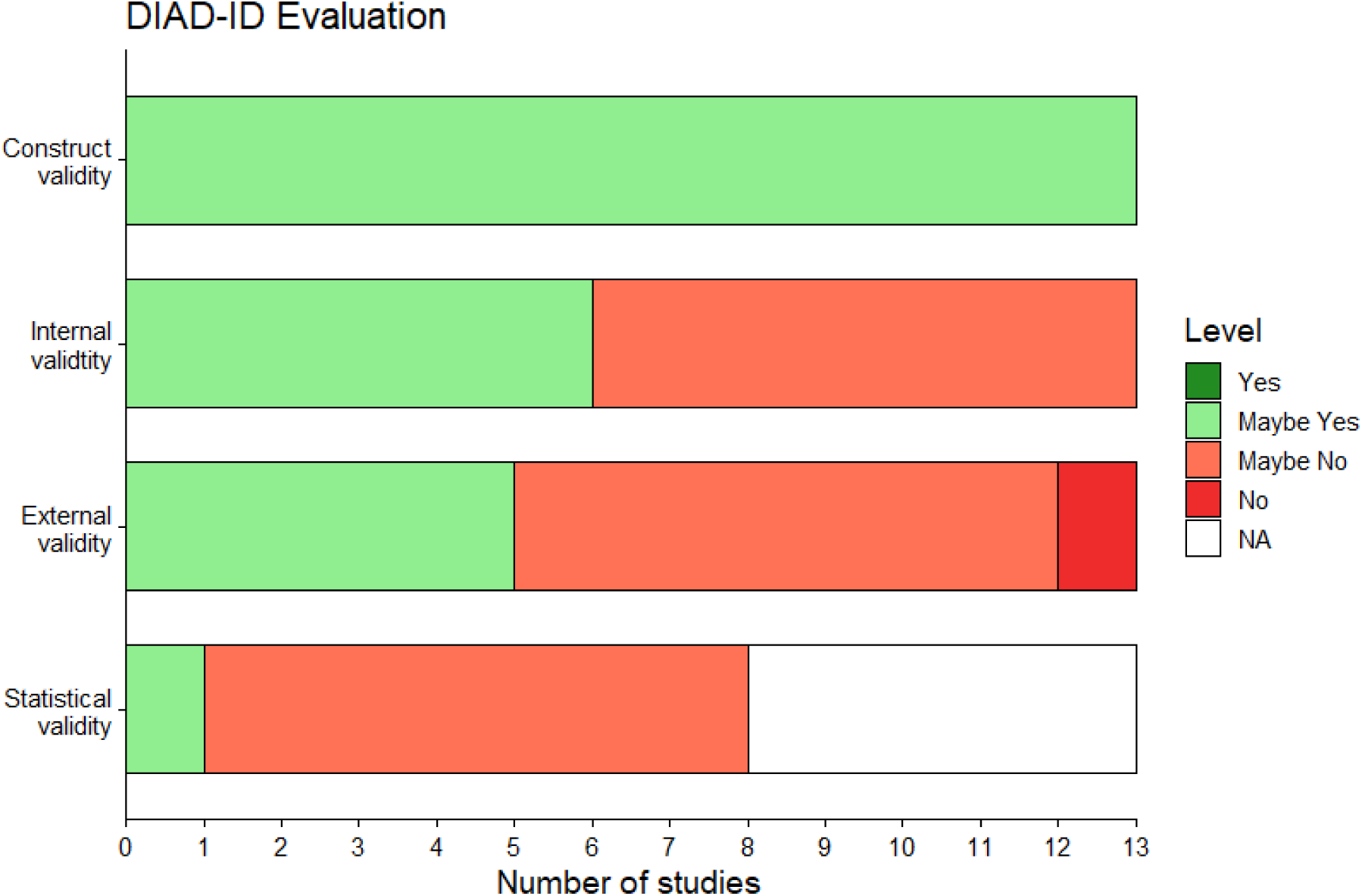
Distribution of DIAD-ID ratings for the four validity criteria. Dark green represents the number of “Yes” ratings (no study received this rating in any category), light green represents “Maybe Yes”, orange represents “Maybe No”, and red represents “No”. Studies that provided data upon request did not receive a rating on statistical validity, hence the white category represents NAs. The y-axis shows the specific validity category, and the x-axis shows the number of studies receiving the rating.

**Figure 5.**
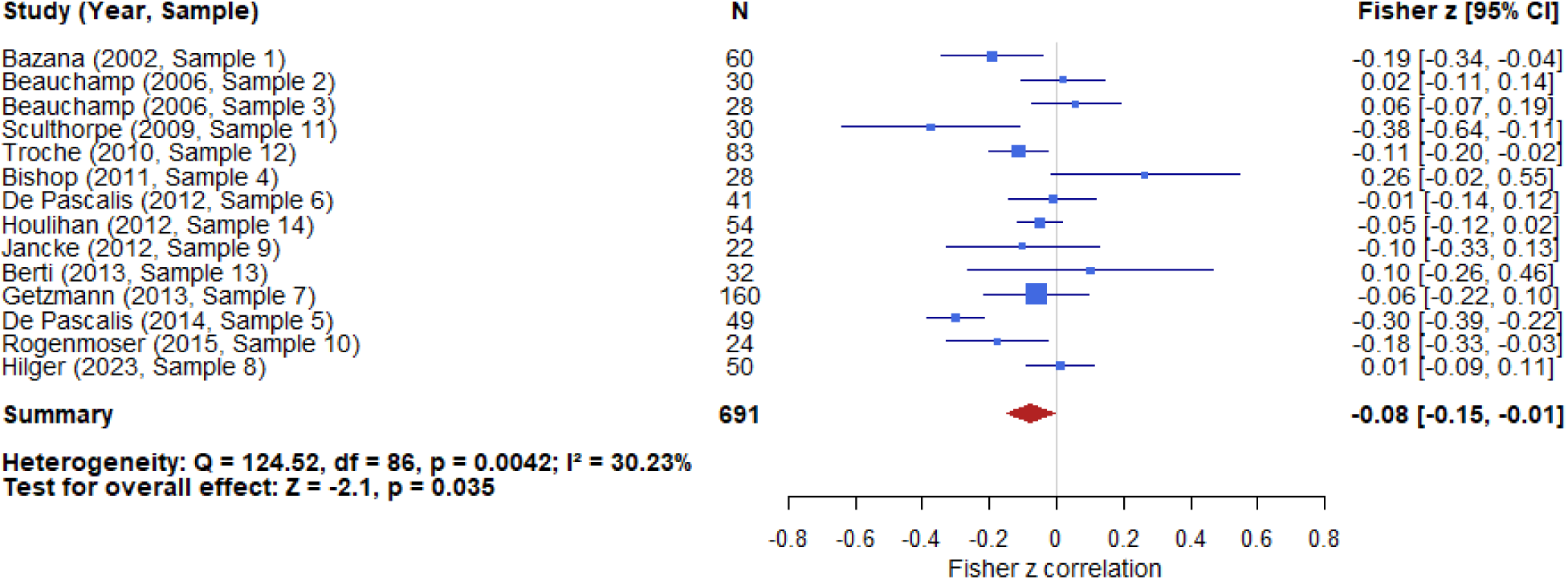
Small, significant association between MMN amplitude and general cognitive ability. Forest plot illustrating study sample-specific correlations between individual variation in MMN amplitude and general cognitive ability (GCA). The blue boxes represent the Fisher’s *z* correlation for the respective study sample, weighted by sample size, with the horizontal lines displaying the 95%-confidence intervals (95% CI). One correlation per sample is depicted, calculated by averaging all relevant sub-effects using inverse variance-weighting. One study contributed multiple samples (Beauchamp & Stelmack, 2006), and most studies reported multiple conditions per sample (occasionally involving subsamples). Box sizes indicate the relative influence of individual samples on the overall meta-analytic estimate, determined by each sample’s weight. The meta-analytically derived overall effect size across samples is depicted by the red diamond. The pooled sample size and Fisher’s *z* with 95% CI are depicted similarly to those for the individual samples. Additionally, the across-sample heterogeneity measures (Q, I²) and the overall effect test statistics (*z*, *p*) are presented in the lower left corner of the figure.

**Figure 6.**
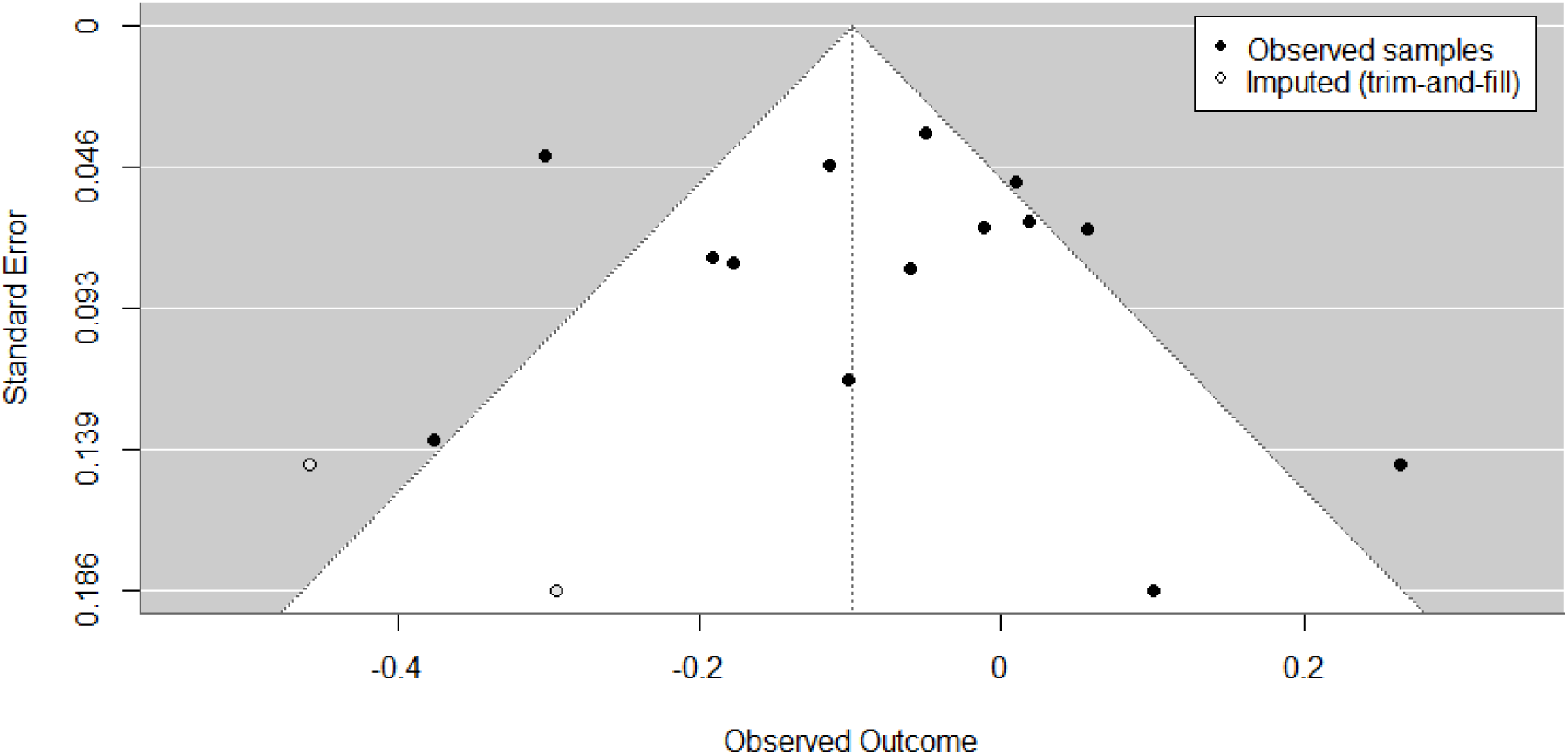
No publication bias in studies on the association between MMN amplitude and general cognitive ability. Funnel plot displaying MMN-GCA amplitude associations (Fisher’s *z*; x-axis) versus standard error (y-axis) with trim-and-fill. Each point reflects one sample; samples with multiple reported effects were combined by applying a fixed-effect inverse-variance weighted mean on the Fisher’s z scale. The two open markers represent the two imputed studies after trim-and-fill.

**Figure 7.**
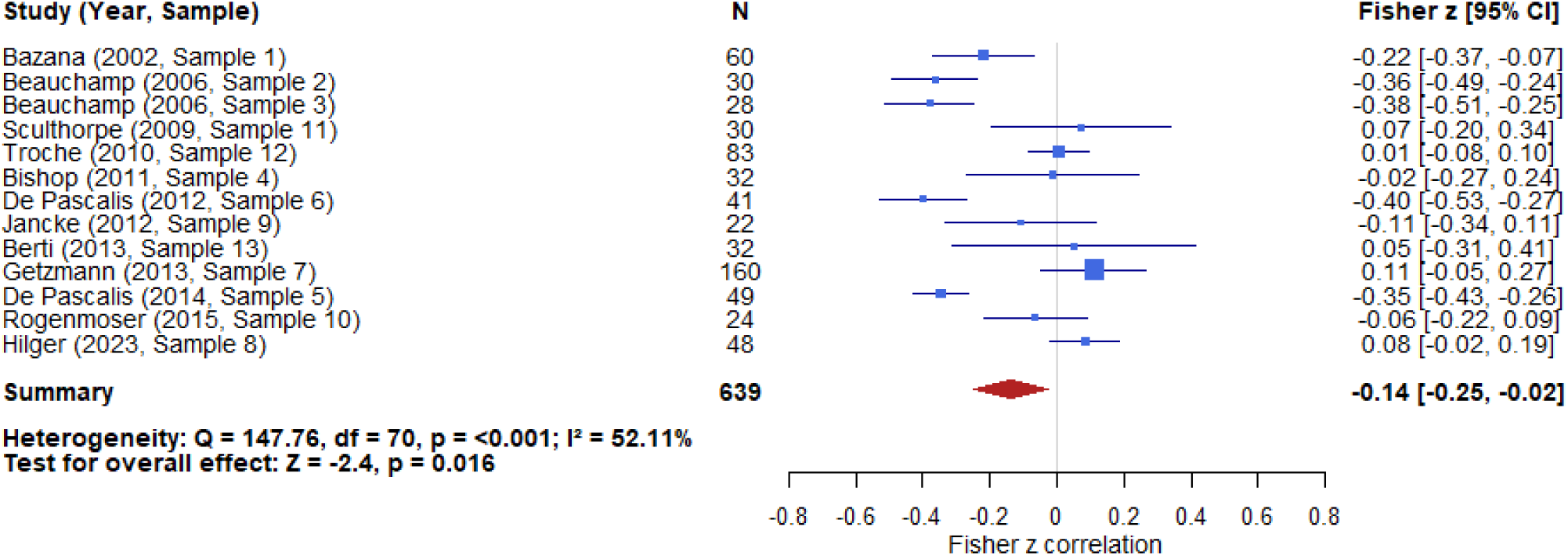
Small but significant association between MMN latency and general cognitive ability. Forest plot showing study sample-specific correlations between individual variation in MMN latency and general cognitive ability (GCA). The blue boxes represent Fisher’s *z* correlation for the respective study sample, weighted by sample site, with the horizontal lines displaying the 95%-confidence intervals (95% CI). One correlation per sample is depicted, calculated by averaging all relevant sub-effects using inverse variance-weighting. One study contributed multiple samples (Beauchamp & Stelmack, 2006), and most studies reported multiple conditions per sample (occasionally involving subsamples). Houlihan and Stelmack (2012) did not report MMN latency and GCA associations. Box sizes indicate the relative influence of individual samples on the overall meta-analytic estimate, determined by each sample’s weight. The meta-analytically derived overall effect size across samples is depicted by the red diamond. The pooled sample size and Fisher’s *z* with 95% CI are depicted similarly to those for the individual samples. Additionally, the across-sample heterogeneity measures (Q, I²) and the overall effect test statistics (*z, p*) are presented in the lower left corner of the figure.

#### 4.2.1 Construct Validity: Does the Study Design Adequately Reflect the Concepts and the Hypotheses?

All 13 included studies showed acceptable levels of construct validity (all 13: “Maybe Yes”). This may be attributable to the well-established operationalization of MMN and intelligence, which facilitates greater comparability and generalizability of measurement processes. However, in most cases, studies did not receive an ideal rating (“Yes” instead of “Maybe Yes”) because of a lack of reliability ratings for MMN latency measures, MMN amplitude measures (questions 1.4.3 and 1.4.4), and for GCA (1.5.4).

#### 4.2.2 Internal Validity: Did the Study Design Allow Clear Conclusions about the Relationship Between MMN and GCA?

The ratings for internal validity revealed more variation, with six studies allowing conclusions to be drawn about the association between MMN and GCA on an adequate level (“Maybe Yes”). In contrast, seven studies did not fully implement a study design that would have permitted clear conclusions regarding these associations (“Maybe No”). While the recording conditions of MMN were often rather well reported and of good methodological quality, most of the “Maybe No” ratings resulted from a lack of information about the GCA assessment (e. g., environment, test administrators, etc.; questions 2.2.1-2.2.5).

#### 4.2.3 External Validity: Do the Samples and Operationalized Constructs Allow the Results to Be Generalized beyond the Study Sample?

Overall external validity ratings were rather low, with only five studies receiving a positive rating (“Maybe Yes”) and eight studies receiving a negative rating (7x “Maybe No”; 1x “No”). While some studies employed GCA measures capturing a broader spectrum of cognitive ability, the majority relied on unidimensional measures. Moreover, due to the applied exclusion criteria, all studies included only healthy participants. Further, the diversity in participant characteristics such as sex and age was limited, as most studies recruited young healthy female students (questions 3.1.2 and 3.2.2).

#### 4.2.4 Statistical Validity: Were Associations Adequately Quantified, and Were Statistical Tests Adequately Reported?

Statistical validity was the category with the lowest rating. While five studies were excluded from this category due to the relevant data being retrieved directly from the authors (rated as “NA”), seven of the other remaining studies were evaluated negatively (“Maybe No”), and only one study received a positive rating (“Maybe Yes”). These low ratings reflect insufficient testing of statistical assumptions or the selection of association measures that were inappropriate given the data (question 4.2.1 and 4.2.2). Moreover, some of the studies did not report statistical tests adequately (questions 4.2.2-4.2.4).

### 4.3 Meta-analytic Synthesis of Amplitude Effects

#### 4.3.1 Hypothesis 1: Negative Association Between MMN Amplitude and General Cognitive Ability

Across samples, MMN amplitudes showed a small but significant association with GCA: *r* = - .077 (87 original study correlations from 14 samples; Fisher’s *z* = −0.078; SE = 0.037; *z* = - 2.104; *p* = 0.03; 95% CI [−0.150, −0.005]). Note that while the association is negative, MMN amplitude is an inherently negative parameter. Thus, a negative association implies that larger absolute amplitude values are positively associated with larger GCA scores. Across-sample heterogeneity was statistically significant (Q(86) = 124.52; *p* = 0.0042), with moderate heterogeneity (I^2^ = 30.23%) and a between-sample variance (σ²) of 0.0126.

#### 4.3.2 Moderator Effects Task Difficulty

Across nine samples, task difficulty was assessed for 59 associations, of which 20 were categorized as elicited during low, another 20 as during medium, and 19 as during high difficulty tasks. The effect of the moderator difficulty was not significant (QM(2) = 2.450; *p* = 0.29), although subsequent intercept analyses indicated the existence of a significant negative association between MMN amplitude and GCA within low difficulty tasks (Fisher’s *z* estimate = −0.17; SE = 0.053; *z*-value = −3.159; *p* = 0.0016; 95% CI [−0.273, −0.064]) but not for the comparison between low and medium and low to high difficulty tasks. Residual heterogeneity remained significant (QE(56) = 83.911; *p* = 0.0093), with between-study variance estimated at σ² = 0.013, and approximate residual I^2^ = 31.5%.

### Age

The moderation effect of mean sample age did not reach statistical significance (QM(1) = 3.455; *p* = 0.06; 69 associations across 11 samples). However, trend-level support for stronger negative correlations between MMN amplitude and GCA in older participants (Fisher’s *z* estimate = 0.015; SE = 0.008; *z*-value = 1.859; *p* = 0.063; 95% CI [-0.001, 0.031]) could be observed. The between-study variance was obtained at σ² = 0.016 with a residual I^2^ = 33.4%, while residual heterogeneity was significant (QE(67) = 98.030; *p* = 0.0080).

### Gender

Gender did not significantly moderate the effect on the association between MMN amplitude and GCA (QM(1) = 1.584; *p* = 0.21; 86 associations across 13 samples). Residual heterogeneity was significant (QE(84) = 121.597; *p* = 0.0046), and the between-study variance was estimated at σ² = 0.015 while the residual I^2^ was computed as 34.1%.

### MMN Amplitude Window Length

The moderator effect MMN mean sample window length was not significant across 13 samples and 83 associations (QM(1) = 0.061; *p* = 0.81). Residual heterogeneity stayed significant (QE(81) = 114.279; *p* = 0.0088), with between-study variance derived at σ² = 0.016, and residual I^2^ was approximately 36.6%.

#### 4.3.3 Examination of Outliers

Two samples (sample 3 from Beauchamp & Stelmack, 2006; De Pascalis et al., 2014) had higher studentized residuals than ± 1.96 (Viechtbauer & Cheung, 2010). However, since neither of those samples demonstrated a Cook’s distance value larger than the median value plus six times the interquartile range (0.522 for the global amplitude model), no sample met the joint criterion for potential outliers. Nevertheless, a sensitivity analysis without these two samples (number of associations post removal: k = 66) reduced the estimated effect slightly (Fisher’s z: −0.078 based on 14 samples vs. −0.064 based on 12 samples), while the *p*-value remained significant (*p* = 0.018) and across-sample heterogeneity became non-significant (Q(65) = 69.633; *p* = 0.325).

#### 4.3.4 Publication Bias

Publication bias was assessed using Egger’s regression test and Begg’s rank correlation test. However, neither test yielded significant results (Egger’s regression: z = 0.722; *p* = 0.47; Begg’s rank correlation: *τ* = 0.011; *p* = 1.00). Thus, these analyses did not indicate evidence of funnel plot asymmetry. Trim-and-fill imputation suggested two missing studies on the left side of the funnel plot (SE = 2.55). Imputing these studies minimally altered the still significant outcome of the meta-analytic model for the association between GCA and MMN amplitude (z = −0.099; *p* = 0.0101; 95% CI [−0.174, −0.024]). Additionally, a fail-safe *N* analysis was performed, suggesting that three unpublished null studies could suffice to render the overall effect non-significant. This is likely due to the relatively small number of studies included, which probably reflects limited robustness rather than the absence of an effect, as a few null findings can disproportionately influence the overall effect size.

### 4.4 Meta-analytic Synthesis of Latency Effects

#### 4.4.1 Hypothesis 2: Negative Association Between MMN Latency and General Cognitive Ability

The across-sample association between MMN Latency and GCA was small but statistically significant: *r* = −.135 (71 original study correlations from 13 samples; Fisher’s *z* = −0.136; SE = 0.057; *z* = −2.402; *p* = 0.016; 95% CI [−0.247, −0.025]). The estimated between-sample variance (σ²) was 0.034, with a significant across-sample heterogeneity (Q(70) = 147.76; *p* < 0.0001) of 52.11% (I^2^).

#### 4.4.2 Moderator Effects Task Difficulty

Eight samples employed tasks with varying difficulty levels, resulting in 16 associations for low and medium difficulty, respectively, and 11 for high difficulty (total number of associations = 43). However, the effect of the moderator task difficulty was not significant (QM(2) = 0.1585; *p* = 0.92). More detailed analyses revealed that the association between MMN latency and GCA was only significant for low difficulty tasks (Fisher’s *z* estimate = −0.16; SE = 0.070; *z*-value = - 2.294; *p* = 0.022; 95% CI [–0.300, −0.024]). Residual heterogeneity was significant (QE(40) = 66.138; *p* = 0.0058), with between-study variance estimated at σ² = 0.024, and approximate residual I^2^ = 44.9%.

### Age

The moderation effect of mean sample age did not reach statistical significance (QM(1) = 0.268; *p* = 0.60; 61 associations across 10 samples). Residual heterogeneity was significant (QE(59) = 100.573; *p* = 0.0006). The between-study variance was derived at σ² = 0.029 while residual I^2^ was approximately at 46.9%.

### Gender

Gender was the only significant moderator of the association between MMN latency and GCA (QM(1) = 8.745; *p* = 0.0031; 62 associations across 11 samples; Fisher’s *z* estimate = −0.96; SE = 0.325; *z*-value = −2.957; *p* = 0.0031; 95% CI [−1.596, −0.324]). This moderating effect was also significant after Bonferroni correction (*p* = 0.0063). In addition, residual heterogeneity did not remain significant (QE(60) = 58.565; *p* = 0.528). The between-study variance was estimated at σ² = 0.010 while the residual I^2^ was computed as 22.9%. These results indicate that for a mean-centered sample, longer MMN latencies are associated with lower GCA scores, while this negative association tends to be stronger in samples with a higher percentage of women. However, these findings should be interpreted with caution, as the percentage of women across the relevant samples was high (mean = 92.5%; median = 100%). Consequently, the limited variance implied reduced power for the moderating effect of gender. Figure 8 displays this issue: The meta-regression prediction is adequately accurate in the 75% to 100% area, where sufficient data were available. However, the prediction accuracy decreases for samples with a lower female proportion. In the area between 50% and 75% (yellow shaded area), only one sample contributed data (Berti et al., 2012). The prediction for lower percentages (<50%) had to be entirely extrapolated (red shaded area). Thus, the implications of the moderation effect should only be generalized to samples with at least 75% women.

**Figure 8.**
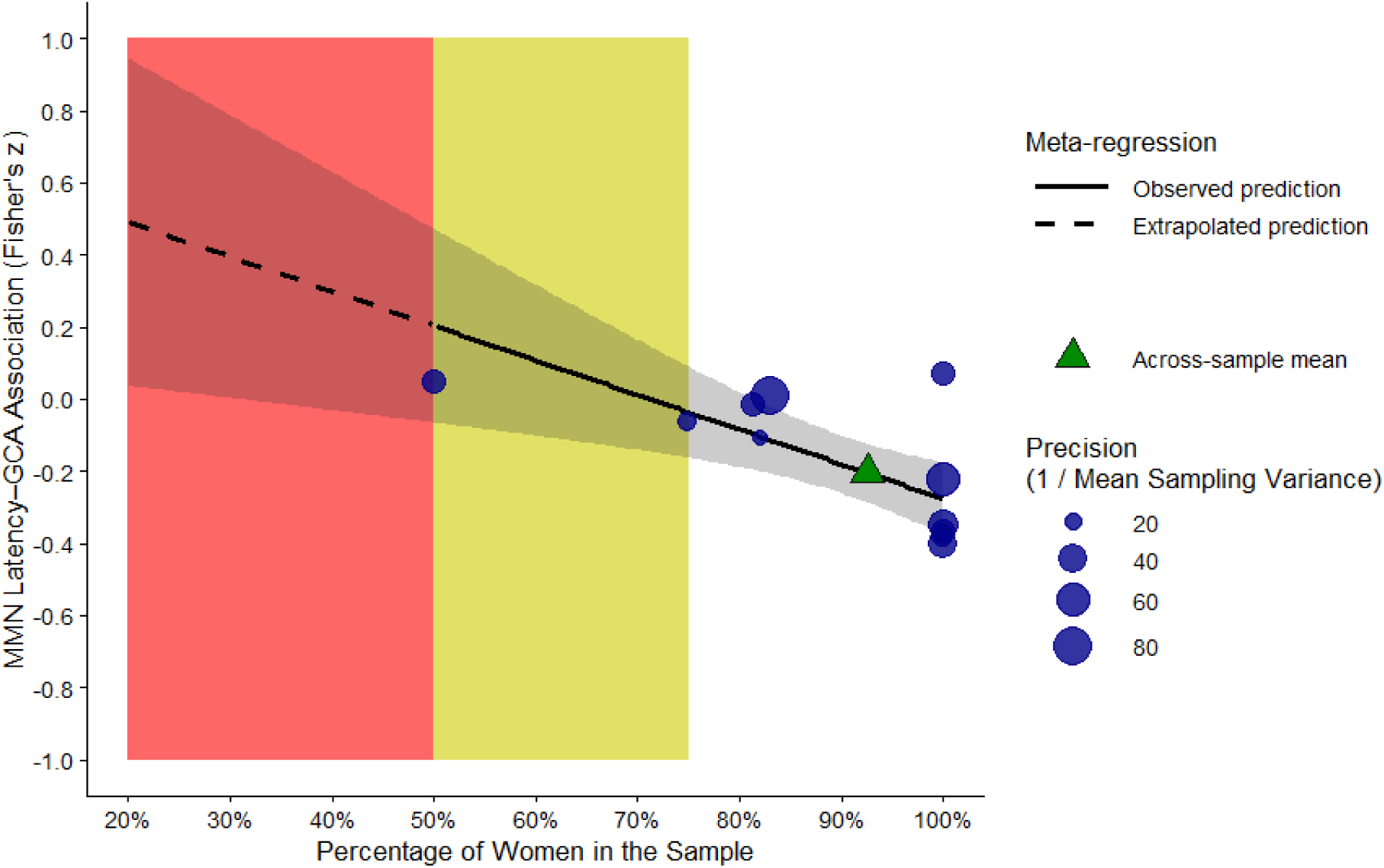
Moderator effect of gender (displayed as percentage female) on the association between MMN latency and general cognitive ability. The solid black line portrays the meta-regression prediction of the influence in the observed area, while the dashed line represents the extrapolated meta-regression prediction. The gray-shaded area around the line represents the 95% confidence interval, with a broader band implying less prediction accuracy. The red shaded area indicates the extrapolated region. The yellow shaded area only contains data from one study. The green triangle marks the across-sample mean percentage of women (92.5%) along with the predicted association between MMN latency and GCA (Fisher’s *z* = −0.20) at that mean. Additionally, the blue dots portray the samples included in this analysis. Their position displays the mean percentage of women in the sample (x-axis) and the mean Fisher’s *z* correlation (y-axis). The size of the circles reflects the relative precision of each sample (1 / mean sampling variance). Thus, samples with larger circles influence the meta-regression more.

**Figure 9.**
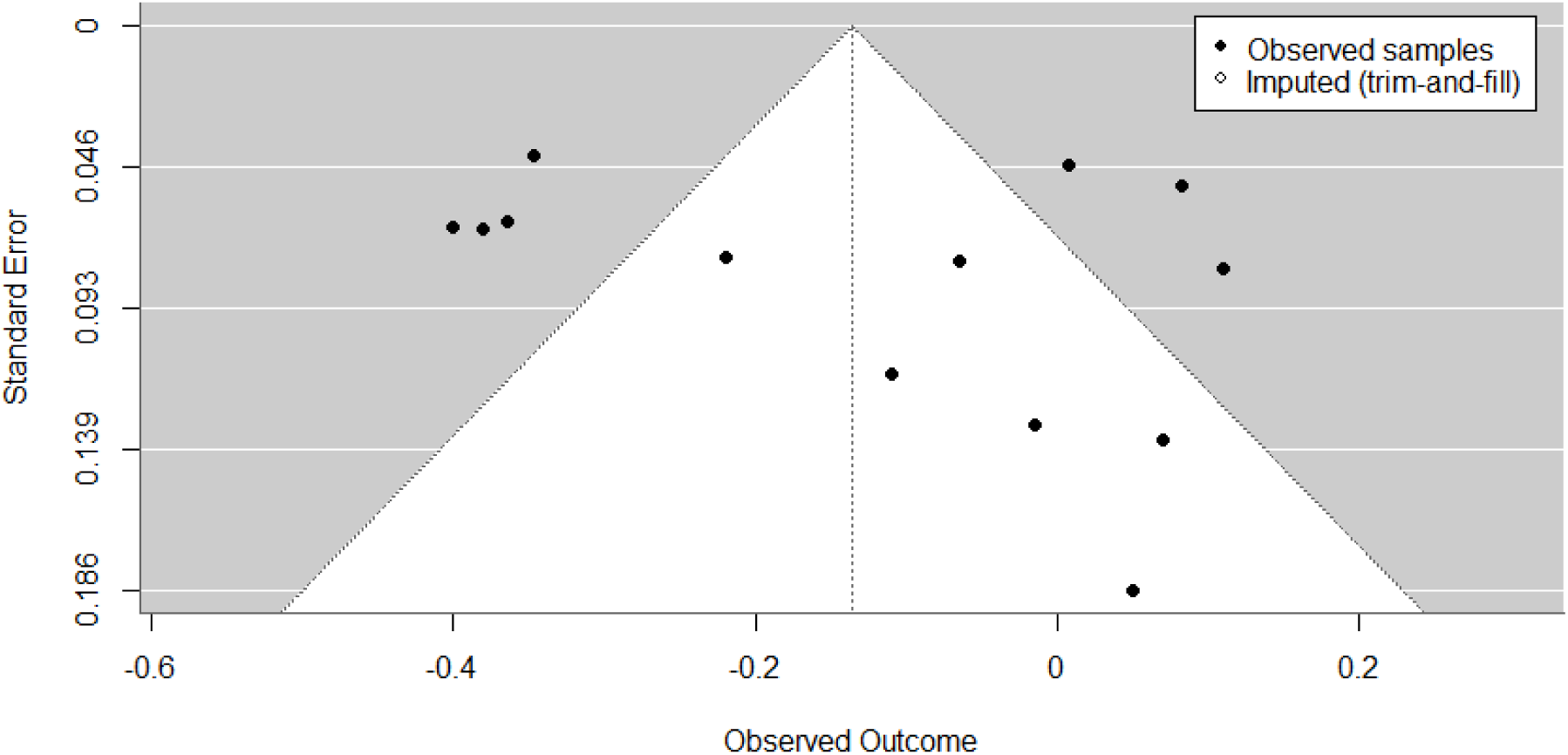
No publication bias in studies on the association between MMN latency and general cognitive ability. Funnel plot displaying MMN-GCA latency associations (Fisher’s *z*; x-axis) versus standard error (y-axis) with trim-and-fill. Each point reflects one study; samples with multiple reported effects were combined by applying a fixed-effect inverse-variance weighted mean on the Fisher’s *z* scale. The trim-and-fill imputation did not indicate any missing studies, which explains the absence of open markers.

### MMN Latency Window Length

The moderator effect for window length was not significant across 12 samples and 67 associations (QM(1) = 0.059; *p* = 0.81). The between-study variance was σ² = 0.039. Residual I^2^ was at 57.0%, while residual heterogeneity stayed significant (QE(65) = 130.041; *p* < .0001).

#### 4.4.3 Examination of Outliers

The joint outlier criterion was not reached by any study. Although one sample (sample 3 from Beauchamp & Stelmack, 2006) exhibited studentized residuals higher than ± 1.96 (Viechtbauer & Cheung, 2010), this sample did not have a Cook’s distance value larger than the median value plus six times the interquartile range (0.830 for the global latency model). Nevertheless, a sensitivity analysis was carried out to assess the influence of the one detected sample (number of associations post removal: k = 62). While the across-sample association was slightly reduced (Fisher’s *z*: −0.136 based on 13 samples vs. −0.114 based on 12 samples), the *p*-value (*p* = 0.047) as well as across-sample heterogeneity (Q(61) = 124.342; *p* < .0001) remained significant.

#### 4.4.4 Publication Bias

The assessment of publication bias did not provide any evidence of funnel plot asymmetry. Neither Egger’s regression test (*z* = 1.353; *p* = 0.176) nor Begg’s rank correlation test (*τ* = 0.231; *p* = 0.306) was statistically significant. The trim-and-fill procedure did not impute any potentially missing studies in the funnel plot, further supporting a low risk of publication bias. A further fail-safe *N* analysis estimated that six unpublished null studies could potentially reduce the overall effect to non-significance. However, this again might be attributable to the relatively small number of studies and should therefore be interpreted with caution.

## 5. Discussion

### 5.1 Summary

This preregistered meta-analysis and systematic review provides the first comprehensive assessment of the current state of research on the association between mismatch negativity (MMN) and general cognitive ability (GCA) in healthy adult participants. 721 individual studies were identified and screened by title and abstract. This resulted in 272 full-texts considered for more detailed assessment. Thirteen studies (comprising 14 samples and 158 effects) were finally included in the qualitative review and meta-analytic comparisons. For the systematic review, study quality and design were evaluated by the Study Implementation Assessment Device for Individual Difference Research (DIAD-ID) (Euler & Hilger, 2026). Corroborating both primary hypotheses, a significant negative across-sample effect emerged for the association between MMN amplitude and GCA (*r* = −.08; 95% CI [−.15, −.01]) as well as for the association between MMN latency and GCA (*r* = −.13; 95% CI [−.25, −.02]). This suggests stronger and faster MMN responses in individuals with higher cognitive ability. Multiple moderators were explored, while only the effect of gender on the MMN latency-GCA association reached significance. Across all analyses, sample heterogeneity was moderate, and no evidence of publication bias was observed. Below, limitations and open questions are discussed, and recommendations for the future of research on electrophysiological correlates of general cognitive ability are presented.

### 5.2 Contextualization of Findings

#### Quality of Literature

Across all studies included in this meta-analysis, the average sample consisted of healthy, young, female participants of above average intelligence (mean age = 24.8, mean percentage female = 88%, sample mean IQ = 111.7). As universities often require participation in research studies for credit, similarly composed samples are common in psychological research. However, the use of these samples critically limits the interpretability of the results and their generalizability to a wider population. Thus, while this meta-analysis yielded significant results, it is important to consider that in such a homogeneous sample, even small effects can pass the significance threshold, and results should only be cautiously applied to groups differing in age and gender distribution.

Further, although all but one study (Houlihan & Stelmack, 2012) reported or provided the association between both MMN components and GCA, descriptive data were documented only sparsely. However, reporting mean MMN amplitudes and latencies, mean Wechsler IQ scores inclusively corresponding score ranges, as well as age-range and mean age for the full sample and all subsamples, is essential not only for transparency and secondary analyses of the data, but also for the interpretation of respective results. For example, descriptive statistics for MMN amplitude and latency were only reported in two out of seven studies (excluding studies that provided the relevant data after mail request). More critically, three studies failed to report the relevant mean ages entirely, either for the full sample or after excluding participants (Berti et al., 2012; Getzmann et al., 2013; Houlihan & Stelmack, 2012), although two of these studies (Berti et al., 2012; Getzmann et al., 2013) had the broadest age range (18-65+).

To systematically assess the quality of all included studies regarding research design, concept implementation, and statistical methodology, the DIAD-ID (Euler & Hilger, 2026) was implemented. All 13 studies showed adequate levels of construct validity. This could mainly be attributed to nearly no study reporting reliability ratings for GCA, MMN amplitude, and MMN latency measures. Eleven studies applied well-established GCA measures like the Multidimensional Aptitude Battery (MAB; Wallbrown et al., 1988), Raven’s (Advanced) Progressive Matrices (RPM; Nurhudaya et al., 2019), or the Mehrfachwahl-Wortschatz-Test (MWT-B; Lehrl et al., 1995), for which reliability estimates are depicted in the manuals. Additionally, all studies were designed at least adequately to elicit MMN. However, reporting reliability ratings calculated within the employed samples would have been important to provide the reader with information about the consistency within individual measurements, with the opportunity to compare measurements across studies and to assess potential attenuation of correlations in this meta-analysis. The evaluation of internal validity yielded mixed results, mostly attributable to a lack of reports of details about the environment and procedures during EEG and GCA recording. Importantly, these ratings do not automatically imply inadequate study design quality but rather highlight the need for more detailed documentation of scientific research. External validity was, on average, rated negatively (5 positive and 8 negative ratings). These negative ratings were primarily driven by sample homogeneity, as discussed above. Lastly, seven out of eight studies considered for statistical validity did not document testing statistical assumptions prior to correlational analysis. However, each of those studies computed Pearson’s product-moment correlations, which should only be applied when data is at least approximately normally distributed (Schober et al., 2018). Thus, without pretesting this assumption, Spearman’s *rho* may have been preferable for reducing variability and increasing robustness (de Winter et al., 2016).

### 5.3 Quantitative Meta-Analysis

#### 5.3.1 Association Between Cognitive Ability and MMN Amplitudes

Each of the 13 included studies provided at least one sample (total number of samples: 14) and one correlation to the across-sample association between MMN amplitudes and GCA. Ultimately, 87 individual correlations across 691 participants contributed to the estimated across-sample effect of −.08 (*p* = 0.035). This small but significant effect supports the first preregistered primary hypotheses (H1), assuming a negative association between MMN amplitude and GCA. The results, therefore, imply a larger neural response (since MMN amplitude is a negative component) to violations of (unconsciously processed) expectations in individuals with higher cognitive abilities. While this result was expected and is plausible, it should be interpreted with caution, especially since the 95% confidence interval nearly includes zero ([−0.150, −0.005]). Nevertheless, this meta-analytically derived finding provides an important baseline estimate of the association between MMN amplitude and GCA for future studies.

### Implications and Interpretation

Since the discovery of MMN in the 1970s, two main theories about its interpretation have emerged: the memory-based hypothesis and the adaptation hypothesis (Fitzgerald & Todd, 2020). The memory-based hypothesis postulates that the MMN is evoked by an automated change-detection mechanism that requires a comparison between the sensory-memory encoding of the antecedent auditory standard stimulus with that of the atypical acoustic input (Näätänen et al., 2005). This theory has been expanded towards the model adjustment hypothesis, proposing that MMN might represent the in-situ adaptation of perceptual representations to unexpected auditory input (Sussman & Winkler, 2001; Winkler et al., 1996). In contrast, the adaptation hypothesis assumes that MMN is not based on a separate process of change detection but rather is an extension of the N1 component (Jääskeläinen, 2004). While the model adjustment hypothesis has historically been predominant, Friston (2005) proposed putting the MMN into a predictive coding framework. This framework combines the other two hypotheses (memory/adjustment vs. adaptation) by integrating the neural adaptations to recurring stimuli into the refinement of the sensory representations after deviants (Garrido et al., 2009). Although the predictive coding framework provides a unifying explanation for MMN generation, it makes no explicit assumptions regarding individual differences and potential associations with GCA. However, it has been argued that small associations between early ERP components like the MMN and GCA might be expected under a hierarchical predictive coding framework, due to the relatively low task and neural demands involved (Euler, 2018).

Furthermore, both MMN amplitude and GCA have been linked to other concepts that may allow to indirectly integrate the MMN amplitude-GCA association into the predictive coding framework. Friston (2005) associates the generation of the MMN with short-term neural plasticity, which is substantiated by studies observing ketamine-induced reductions in synaptic plasticity leading to reduced MMN amplitudes (Schmidt et al., 2012). Furthermore, Thatcher et al. (2016) propose that the mechanism of homeostatic neuroplasticity supports more efficient information flow in higher IQ individuals. The results of this meta-analysis, thus, could be considered consistent with the predictive coding framework and might also help to understand the connection between MMN amplitude, GCA, and short-term neural plasticity by adding a small association between MMN amplitude and GCA. However, the potential associations between these three factors, though plausible, remain hypothetical and therefore require more empirical research to validate the theoretical assumptions.

Moreover, the predictive coding framework postulates that the MMN is elicited once an individual fails to predict sensory input via a generative model (pre-trained by a frequent stimulus) and consequently fails to minimize prediction errors (PE; Garrido et al., 2009). The meta-analytic findings could thus be interpreted as individuals with higher cognitive ability showing larger negative deflections following PEs, while the connection between stronger responses to PE and GCA could potentially be explained by PE playing a big role in sensorial learning and memory encoding (Henson & Gagnepain, 2010). This theory was supported by Greve et al. (2017), who found improved memory after larger PE, caused by a situation where the standard stimulus combination was more frequently shown than a different combination of visual stimuli, as well as by Giard et al. (1990), who linked the primary MMN generator, the superior temporal gyrus, to a sensory mechanism comparing perceived stimuli to preformed cognitive representations. Finally, the contribution of both bottom-up (e.g., sensorial encoding) and top-down processes (e.g., prediction of future stimuli), as well as proposals about the existence of “primitive sensory intelligence” (Näätänen et al., 2007), based on involved automatic cognitive mechanisms such as early encoding of stimuli and attributes (Saarinen et al., 1992) and unattended rule registration and assessment (Paavilainen et al., 2007), support the link between MMN amplitude and GCA further. Based on these considerations, it could be speculated that mismatch negativity amplitude relates to general cognitive ability, because larger MMN amplitudes reflect enhanced sensorial encoding and learning concerning the prediction of stimuli. This improved ability to predict might depend on larger prediction errors, which would ultimately result in better generative models in higher cognitive ability individuals. However, improving these generative models is not expected to eliminate MMN entirely. Instead, while it could be hypothesized that model adaptation would initially reduce MMN after the first few presentations of deviant stimuli, an ideal prediction model would still predict the more probable standard stimulus instead of the deviant. These theoretical assumptions, however, require future empirical testing.

Furthermore, while the overall association between MMN amplitude and GCA reached statistical significance, the small effect size (*r* = −.08) necessitates cautious interpretation. In addition, even though a previous article by Wacongne et al. (2015) provides a neurobiological model that connects MMN generation and predictive coding with NMDA receptors, the framework has also faced criticism. For example, predictive coding has been criticized for being a more theoretical than applicable conceptualization compared to the adaptation model (May, 2021). Therefore, to gain a better understanding of the association between MMN amplitude and GCA as well as its theoretical conceptualization, adequate operationalization is essential.

### Moderator Effects

None of the tested moderators revealed a significant effect on the association between MMN amplitude and GCA. Although task difficulty has frequently been manipulated in previous ERP research (e.g., Ghani et al., 2020), the explicit operationalizations vary considerably. For instance, manipulating the difficulty of the active task, while the passive component of the task elicits MMN, has only been weakly associated with MMN amplitude or not at all (Dittmann-Balcar et al., 1999; Haroush et al., 2010; Muller-Gass et al., 2006). Task difficulty in the present meta-analysis was conceptualized as the difficulty to discriminate between standard and deviant stimuli or patterns in the MMN eliciting paradigm (see 3.3, “Task Difficulty”). Stimuli with higher discrimination difficulty are generally associated with eliciting smaller MMN deflections (Amenedo & Escera, 2000; Näätänen et al., 1989). However, since moderation analyses did not yield any significant effects, further research examining discrimination difficulty between stimuli and its impact on the association between GCA and MMN amplitude is required. Concerning the other moderators, for instance, age and gender, it should be noted that the variability was limited, which may have affected the likelihood of detecting an effect.

#### 5.3.2 Association Between Cognitive Ability and MMN Latency

The second hypothesis (H2) postulated the existence of a significant negative association between MMN latencies and GCA. This was supported by the meta-analytic results. Across the 13 included samples, including 639 participants (71 associations in total), the estimated across-sample association between MMN latency and GCA was −0.13 (*p* = 0.016), suggesting faster MMN generation in individuals with higher cognitive ability. However, these findings should also be interpreted conservatively, as the 95% CI is close to zero (95% CI [−0.247, − 0.025]) and the estimated correlation is small. Nonetheless, this meta-analysis provides the first comprehensive overview of the association between MMN latency and GCA and again sets an important baseline estimate for future research.

### Implications and Interpretation

As discussed above, in the light of the predictive coding framework, the MMN could be seen as a representation of prediction errors, caused by a mismatch between the expected and the actual sensory input (Garrido et al., 2009). Based on the findings of this meta-analysis, it could thus be speculated that these comparison processes happen faster in individuals with higher GCA (as they show smaller MMN latencies). This is in line with the results of De Pascalis et al. (2014), who attributed these benefits of mental speed to optimized and therefore accelerated neural signaling without making any concrete statements about what such accelerated neural signaling consists of.

Irrespective of the proposed predictive coding hypotheses, the association between cognitive ability and mental speed operationalized through ERP component latencies (Amin et al., 2015; Fjell & Walhovd, 2003; Schubert et al., 2022) has been widely discussed, with empirical research yielding heterogeneous results: Some studies claimed moderate to strong associations between GCA and neural processing speed, which would be supported by this meta-analysis, while other studies did not find associations (Ociepka et al., 2022; Posthuma et al., 2001). Additionally, Schubert et al. (2018) showed in an experimental design that nicotine-enhanced cognitive processing speed does not improve participants’ results on intelligence tests. The authors proposed that attentional components and salience-network-related processes might moderate the connection between faster cognitive processing and higher cognitive abilities. However, the MMN is automatically elicited, largely independent of controlled attention (Kathmann et al., 1999; Sussman et al., 2003). Nevertheless, the concept of salience network efficiency being a modulating component of processing speed and intelligence (Hilger et al., 2017; Schubert et al., 2018; Töllner et al., 2011) might be supported by the findings of this meta-analysis. Non-auditory MMN generators may be conceptually linked to salience assessment in perception (Todd et al, 2012), and MMN is widely recognized as a sensory deviant-detection mechanism (Näätänen & Alho, 1995). Therefore, it could be assumed that saliency and the associated network play an important role in MMN elicitation. For instance, higher saliency contributes to faster stimulus detection in visual tasks (Nothdurft, 1993) as well as to higher fixation probability, even in peripheral, not actively attended objects (Peacock et al., 2023). Even though these concepts have mainly been researched in visual perception, Huang and Elhilali (2017) found evidence that increased auditory salience also enhanced reaction time. Hence, by transferring these visual concepts to passive auditory detection, it could be hypothesized that more salient stimuli are detected with a higher probability. Consequently, this theoretically implies that highly salient deviants should be detected more efficiently and thus faster. Additionally, a more efficient saliency network, therefore, should hypothetically also enhance deviant-detection speed.

In conclusion, given the small across-sample association (*r* = −.13), mismatch negativity latency alone is unlikely to serve as a neurobiological marker of general cognitive ability with high direct practical relevance. However, the association might serve as an indicator of underlying components and mechanisms, such as faster comparison between prediction and input or more efficient saliency networks, which might modulate MMN latency (and therefore processing speed) as well as GCA.

### Moderator Effects

Only one of the four tested moderators revealed a significant effect, namely gender. Potential reasons for these diverging findings can be considered. First, the analysis of the MMN window length was only exploratory and unlikely to yield any significant results. Regarding age, the absence of any moderation effect could be explained by the meta-analysis only including a few individuals potentially at risk of age-induced cognitive decline (Miller et al., 2009), leading to an artificially reduced variation in cognitive ability at an older age. Additionally, age does not seem to significantly influence MMN latency (Criel et al., 2023). For task difficulty, the lack of a moderation effect could also be based on the operationalization employed in this meta-analysis, as differences in task difficulty across studies could not be considered. However, gender had a large influence on the association between MMN latency and GCA, indicating that stronger negative correlations were present in samples with more women. Although this effect seems to be rather substantial, it should only be interpreted for studies with at least 75% women, as most of the included studies used primarily female samples. Gender differences regarding the general lateralization of MMN processing (Ikezawa et al., 2008; Voyer & Flight, 2001) as well as heterogeneous findings about the influence of gender on MMN latencies (Criel et al, 2023; Kasai et al., 2002; Matsubayashi et al., 2008; Schwade et al., 2017; Toufan et al., 2021) have been previously reported. However, these findings do not explain the moderation effect found in this meta-analysis. In addition, GCA – quantified in a similar way as in this meta-analysis – does not significantly differ between genders (Daseking et al., 2017; Giofre et al., 2022). Therefore, more research is required to explore why higher cognitive abilities in women might be more strongly related to faster latencies than in men.

### 5.4 Limitations

There are limitations to this meta-analysis. First, this meta-analysis includes only 13 studies. The number of studies was limited by the focus on healthy adult samples to increase generalizability. This meant that many studies were already excluded in the title and abstract screening phase due to underage samples or clinical focus. Additionally, 196 of the 256 full-text screened studies either only measure GCA or MMN or were classified as gray literature (for details, see Figure 1). However, as this meta-analysis is the first to address the relation between MMN and GCA at all, the aim was to generate results as generalizable as possible and to provide correlative estimates that can serve as a lower limit to power calculations in future studies. More specific meta-analyses (e.g., based on the relationship between MMN and GCA in dementia patients) should be considered as an important subject for the future. Another limitation refers to the decision not to consider *p*-values for the assessed associations. Although these scores were seldom reported in sufficient detail, implementing significance as an additional criterion can provide further insights, also about common reporting practices and potential publication bias. Finally, Spearman’s *rho* was used to calculate MMN and GCA associations for the provided data. While this is more robust to violations of the normality assumption than Person’s *r* (de Winter et al., 2016), visual inspection of the data implied that monotonicity could not always be unequivocally assumed. Additionally, other limitations, such as sample homogeneity, were addressed in prior parts of the discussion.

### 5.5 Future Directions

This meta-analysis discloses multiple pressing questions and promising further developments that should be addressed in future research. First, the results of the DIAD-ID reveal room for improvement regarding study design quality. Most critically, included studies lacked the appropriate reporting of sample statistics (e.g., reporting participant descriptives after exclusion and for each subsample) and of details regarding the statistical procedure (e.g., whether assumptions of statistical procedures were tested). Thus, developments in this field towards better reporting practices are mandatorily required and can be supported by recent recommendations (Euler & Hilger, 2026).

Second, this meta-analysis points towards various open research opportunities. For instance, the MMN has been explored as a predictor of cognitive changes associated with deterioration of mental health, especially in at-risk individuals (Erickson et al., 2016; Näätänen et al., 2016; Salisbury et al., 2002). This reflects a very important part of previous and future MMN research. Furthermore, it is also essential to understand the underlying mechanisms and cognitive components linked to MMN in healthy adults. The predictive coding framework provides a theoretical foundation for the development of future studies to close this gap. For example, exploring the influence of varying prediction error sizes on the association between MMN and GCA, in line with the predictive coding framework, could prove crucial for testing the hypothesized connections made in this meta-analysis.

Lastly, stimuli varying in saliency could be used to assess the speculated impact of salience network efficiency on MMN latency and GCA. It should be noted that, since these hypotheses are speculative, it cannot be ruled out that these future directions might not find associations between the domains of interest. However, that could potentially point towards other mechanisms modulating the connection between MMN and GCA.

Consequently, the findings of this meta-analysis encourage research on more heterogeneous samples (gender-balanced and across lifespan) while also exploring the underlying mechanisms of the MMN and GCA association.

### 5.6 Conclusion

In conclusion, the results of this meta-analysis and qualitative review suggest that higher intelligence is weakly but significantly associated with larger MMN amplitudes and shorter MMN latencies. Additionally, the association between GCA and MMN latencies appears to be stronger in women. However, although these findings were significant, the small effect sizes, homogeneous samples, and limited number of included studies warrant cautious interpretation. As highlighted in the systematic review, while MMN and GCA were predominantly operationalized and measured in an adequate manner, generalizability and statistical reporting should be optimized across future studies. Consequently, the results of this meta-analysis contribute to positioning the MMN in the context of GCA research but also reveal the need for deeper research on the underlying mechanism, more heterogeneous samples, and more detailed documentation of descriptive data and statistical procedures. By addressing these concerns, future research should further explore the suitability of the MMN event-related brain potential as a relatively easily assessable marker for neural processes implicated in human intelligence and general cognitive ability.

## 6. Author statements

### Author contributions

**Tobias Nöth:** Data curation, Formal analysis, Methodology, Project administration, Writing - Original Draft, Visualization. **Matt Euler:** Formal analysis, Methodology, Writing – Review and Editing. **Kirsten Hilger:** Conceptualization, Data curation, Funding acquisition, Methodology, Project administration, Supervision, Writing – Original Draft.

## Declaration of generative AI and AI-assisted technologies in the manuscript preparation process

During the preparation of this work the authors used ChatGPT in order to assist with writing code. After using this tool, the authors reviewed and edited the content as needed and take full responsibility for the content of the published article.

## Acknowledgements

Our sincere appreciation goes to all authors that provided data.

